# Demultiplexing the heterogeneous conformational ensembles of intrinsically disordered proteins into structurally similar clusters

**DOI:** 10.1101/2022.11.11.516231

**Authors:** Rajeswari Appadurai, Jaya Krishna Koneru, Massimiliano Bonomi, Paul Robustelli, Anand Srivastava

## Abstract

Intrinsically disordered proteins (IDPs) populate a range of conformations that are best described by a heterogeneous ensemble. Grouping an IDP ensemble into “structurally similar” clusters for visualization, interpretation, and analysis purposes is a much-desired but formidable task as the conformational space of IDPs is inherently high-dimensional and reduction techniques often result in ambiguous classifications. Here, we employ the t-distributed stochastic neighbor embedding (t-SNE) technique to generate homogeneous clusters of IDP conformations from the full heterogeneous ensemble. We illustrate the utility of t-SNE by clustering conformations of two disordered proteins, A*β*42, and a C-terminal fragment of *α*-synuclein, in their APO states and when bound to small molecule ligands. Our results shed light on ordered sub-states within disordered ensembles and provide structural and mechanistic insights into binding modes that confer specificity and affinity in IDP ligand binding. t-SNE projections preserve the local neighborhood information and provide interpretable visualizations of the conformational heterogeneity within each ensemble and enable the quantification of cluster populations and their relative shifts upon ligand binding. Our approach provides a new framework for detailed investigations of the thermodynamics and kinetics of IDP ligand binding and will aid rational drug design for IDPs.

**Significance:** Grouping heterogeneous conformations of IDPs into “structurally similar” clusters facilitates a clearer understanding of the properties of IDP conformational ensembles and provides insights into ”structural ensemble: function” relationships. In this work, we provide a unique approach for clustering IDP ensembles efficiently using a non-linear dimensionality reduction method, t-distributed stochastic neighbor embedding (t-SNE), to create clusters with structurally similar IDP conformations. We show how this can be used for meaningful biophysical analyses such as understanding the binding mechanisms of IDPs such as *α*-synuclein and Amyloid *β*42 with small drug molecules.

**Graphical Abstract:** 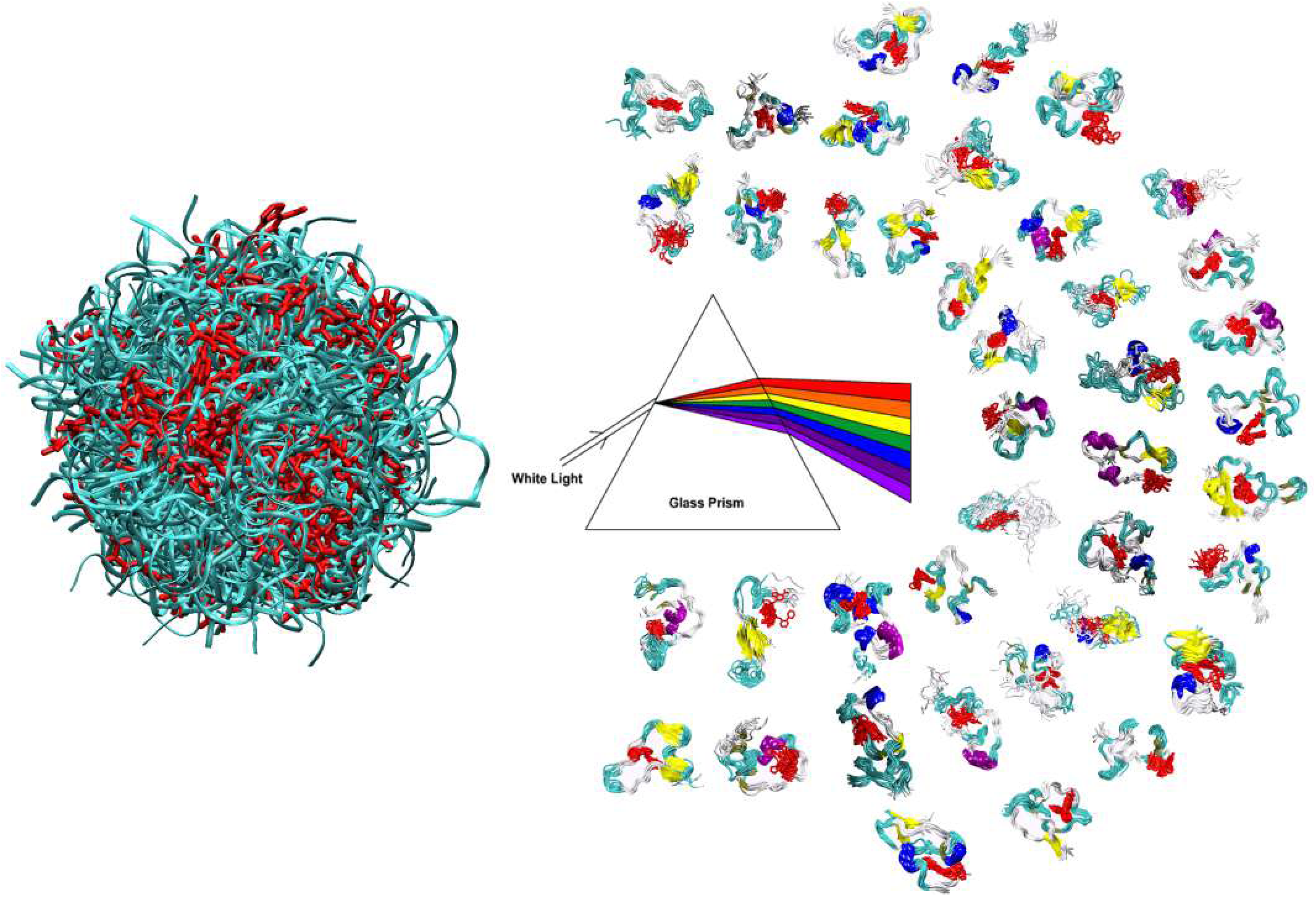

## I. Introduction

In general, knowledge of the 3-dimensional structure of a protein is the first step toward a molecular-level mechanistic understanding of its biological function. This knowledge is also central to activities such as the rational design of drugs, inhibitors, and vaccines and in the broad area of protein engineering and biomolecular recognition^1–7^. With the advances made in structure determination techniques^8–16^ and recent transformative leaps made in computationally predicting the structure from sequence^17–19^, the science of structural biology is going through paradigmatic changes where the knowledge of structure is not the biggest bottleneck anymore^20^. However, outside the realm of these structured proteins exist a ”dark” proteome of intrinsically disordered proteins (IDPs) that constitute more than 40% of all known proteins and play important roles in cellular physiology and diseases^21–27^. An IDP can populate a heterogeneous ensemble of conformations and is functional without taking a unique structure. In essence, IDPs are expanding the classical hypothesis of sequence-structure-function to the sequence-disordered ensemble-function(s) paradigm. Though solution-based experiments like NMR, FRET, and SAXS do provide structural information for IDPs, they generally report time and ensemble-averaged properties of IDP conformations^28–30^. In the absence of computational models, solution experiments are challenging to interpret in terms of individual atomic resolution structures that constitute IDP ensembles. In other words, IDPs are not directly amenable to conventional high-resolution structure determination, structure-based functional correlation, protein engineering, and drug-designing strategies that hinge upon the knowledge of a reference 3-dimensional structure.

Computational tools, particularly those that incorporate the available experimental information, can be effectively used to generate high-resolution ensemble structures of IDPs. Of late, several broad classes of different approaches have been developed for this purpose. Methods based on pre-existing random coil library and simple volume exclusions (examples: Flexible Meccano^31^, TraDES^32^, BEGR^33^) are often used to create an initial exhaustive pool of conformations, which are further processed to produce refined ensembles upon combining with experimental constraints^30,34–39^. These methods, though purely statistical in nature, provide a computationally efficient approach to calculating IDP conformational ensembles that are consistent with experimental data. The second set of approaches utilizes Physics-based molecular simulations either in a coarse-grained representation (examples: SIRAH^40^, ABSINTH^41^, AWSEM-IDP^42^, SOP-IDP^43^, HPS^44^ and others) or with an all-atom resolution,^45–48^ to generate initial Boltzmann-weighted conformational ensembles that can be further refined with experimental restraints using various reweighing approaches^49–51^. Recently developed molecular mechanics force fields for IDPs^45–48^ used in combination with parallel tempering based enhanced sampling approaches such as Replica exchange solute tempering (REST)^52–55^ and hybrid tempering (REHT)^56^ has also shown promise in producing atomic-resolution accurate IDP ensembles consistent with experimental solution data without any added bias in the simulations.

While significant advances have been made in generating high-resolution IDP conformational ensembles that are consistent with experimental data, the subsequent interpretation of these ensembles to address key biological questions related to the interactions of IDPs remains extremely challenging. IDP conformational ensembles are inherently extremely high-dimensional. That is, the phase space of IDPs consists of several thousands of features, which may vary relatively independently, making it extremely challenging to uncover correlations in conformational features among conformations contained in IDP ensembles. This often makes sequence-ensemble-function relationships of IDPs very difficult to understand, even when aided by relatively accurate IDP conformational ensembles. If one could efficiently identify representative conformational sub-states in IDP ensembles, and quantify their relative populations in different molecular and cellular contexts, it would become significantly easier to identify conformational features of IDPs that may be associated with specific functional roles or disease states^57–59^. Therefore, parsing the heterogeneous ensemble data into representative conformational states can be as critical as the generation of the ensemble itself as it allows one to leverage conventional structural-biology analysis tools for IDPs.

The process of dividing large abstract data set into a number of subsets (or groups) based on certain common relations such that the data points within a group are more similar to each other and the points belonging to different groups are dissimilar is called clustering. Due to its ability to provide better visualization and statistical insights, clustering is ubiquitous in the analyses of big-data biological systems with wide-ranging applications such as profiling gene expression pattern^60,61^, de novo structure prediction of proteins^62,63^, the quantitative structure-activity relationship of chemical entities^64^, docking and binding geometry scoring^65^, and also in analyses of protein ensemble from molecular dynamics (MD) trajectory^66^. However, the clustering of IDP ensembles is formidable owing to their large conformational heterogeneity and often different conformations of IDP have similar projected collective variables (CVs). To illustrate this, we present a set of conformations from a simulated IDP ensemble with the same value of R*_g_* as a CV (Fig. S1 in Supplementary Material (SM)). It is evident from this illustration how this could lead to ambiguous classification.

Theoretically well-grounded dimensionality reduction (DR) techniques are now commonly being used in protein conformation analysis to extract the latent low dimensional features and the quantum of information lost during the projection depends heavily on the kind of data set under consideration^67–72^. For example, a highly heterogeneous data set that lies on a high-dimensional manifold as in the case of IDPs is best handled with the non-linear dimension reduction (NLDR) techniques, which generally attempt to keep the nearest neighbors close together. While methods such as ISOMAP and Local Linear Embedding are best suited to unroll or unfold a single continuous manifold, the recently developed t-Distributed Stochastic Neighbor Embedding (t-SNE) method may be more suitable for clustering IDP conformations as it helps to disentangle multiple manifolds in the high-dimensional data concurrently by focusing on the local structure of the data to extract clustered local groups of samples. Consequently, t-SNE tends to perform better in separating clusters and avoiding crowding. Here, we show that t-SNE is particularly well-suited for clustering seemingly disparate IDPs conformations into homogeneous subgroups since it is designed to conserve the local neighborhood when reducing the dimension, which ensures similar data points remain equivalently similar and dissimilar data points remain equivalently dissimilar in the low dimensional and high dimensional space^73^. Due to its ability to provide a very informative visualization of heterogeneity in the data, t-SNE is being increasingly employed in several applications such as clustering data from single cell transcriptomics^74–77^, mass spectrometry imaging^78^, and mass cytometry^79,80^. Lately, t-SNE has also been used for depicting the MD trajectories of folded proteins^81–87^ and for interpretation of mass-spectrometry based experimental data on IDPs by juxtaposing with classical GROMOS-based conformation clusters from the corresponding molecular simulation trajectories of the IDP under consideration^88^.

In this paper, we demonstrate the effectiveness of t-SNE (in combination with K-means clustering) for identifying and visualizing representative conformational substates in IDP ensembles. We investigate the small molecule binding properties of Amyloid *β*42 (A*β*42) and *α*-synuclein (*α*S), proteins involved in the neurodegenerative proteinopathies like Alzheimer’s and Parkinson’s diseases, respectively. Therapeutic interventions by sequestering the monomeric state of these IDPs have recently been explored using state-of-the-art biophysical experiments and long timescale molecular simulations^89,90^. A set of repurposed small molecules such as the c-Myc inhibitor-G5 (benzofurazan N-([1,1-biphenyl]-2-yl)-7-nitrobenzo[c][1,2,5]oxadiazol-4-amine (10074-G5)) and a Rho kinase inhibitor - Fasudil (along with the high-affinity Fasudil variant Ligand-47) have been identified as promising agents against the monomers of A*β*42 and *α*S, respectively. Since the monomeric states of these IDPs are extremely heterogeneous, it is not fully understood how the different conformations form viable complexes with these small molecules and what molecular features derive their affinity and specificity. This insight is obscured by inefficient clustering of the IDP structures using the classical clustering tools. Here we revisit the molecular trajectories of A*β*42 (a total of 56 *µ*secs) and *α*S (total of 573 *µ*secs) using t-SNE (in combination with K-Means clustering). This exercise has improved our knowledge of the binding mechanism of small molecules to such IDPs and also provides us with strategies for designing specific inhibitors with high-affinity binding. Additionally, our clustering analysis provides valuable insight for understanding the conformational landscape of APO and ligand-bound IDPs, which are otherwise hard to obtain. We believe that the method presented here is general in nature and can be used to cluster and visualize IDP ensembles across systems with varying degrees of structural heterogeneity and assist in detailed structural, thermodynamics, and kinetics analyses of IDP conformations in APO and bound states.

## II. Results and Discussion

We aim to cluster the heterogeneous mixture of disordered protein conformations into a subset of unique and homogeneous conformations. To do this, as a first step, we employ t-SNE that projects the large dimensional data in lower dimensions. We then apply K-means clustering on the projections to identify the clusters in the reduced space. Before we illustrate the power of this algorithm as a faithful clustering tool for realistic IDP ensembles, we use a simple alanine-dipeptide (ADP) toy model to provide physical intuition into how t-SNE works. Please see Fig. S2 and the subsection titled *”Physical intuition into t-SNE-based clustering algorithm using alanine dipeptide”* in SM. We use this model system to introduce the role of the critical hyper-parameter perplexity in the t-SNE algorithm, and prescribe a strategy to determine its optimal value for effective clustering. We then apply the method for the analyses of IDP ensembles of complex systems such as A*β*42 and *α*S, each in the presence and absence of small-molecule inhibitors. We list all the systems under consideration in Table I below. We represent the conformations within A*β*42 and *α*S ensembles by the inter-residue Lennard-Jones contact energies and the Cartesian coordinates of heavy atoms, respectively. These measures were chosen for consistency with previous analyses performed on these trajectories^89,91^ to enable faithful comparisons. t-SNE was performed based on the pairwise RMSD of Lennard-Jones contact energies among conformations of A*β*42, and the pairwise backbone RMSDs among conformations of *α*S.

**TABLE I:**
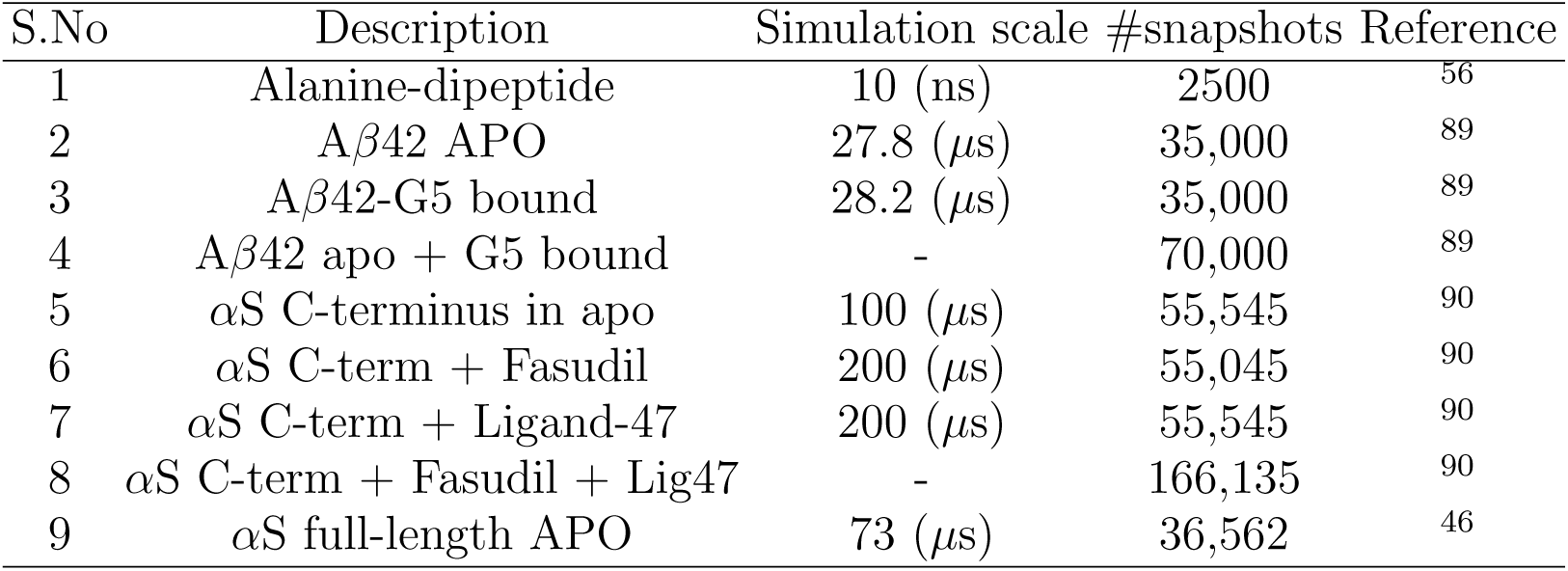
Information on systems and trajectories used in this study

| S.No | Description | Simulation scale | #snapshots | Reference |
| --- | --- | --- | --- | --- |
| 1 | Alanine-dipeptide | 10 (ns) | 2500 | 56 |
| 2 | Aβ42 APO | 27.8 (μs) | 35,000 | 89 |
| 3 | Aβ42-G5 bound | 28.2 (μs) | 35,000 | 89 |
| 4 | Aβ42 apo + G5 bound | - | 70,000 | 89 |
| 5 | αS C-terminus in apo | 100 (μs) | 55,545 | 90 |
| 6 | αS C-term + Fasudil | 200 (μs) | 55,045 | 90 |
| 7 | αS C-term + Ligand-47 | 200 (μs) | 55,545 | 90 |
| 8 | αS C-term + Fasudil + Lig47 | - | 166,135 | 90 |
| 9 | αS full-length APO | 73 (μs) | 36,562 | 46 |

### A. Prescription for choosing optimal parameters for t-SNE clustering of IDPs

The results of t-SNE depend largely on the choice of perplexity. Since the objective criterion here is to maximize clustering, we adopt the well-known Silhouette score,^92^ commonly used for optimizing the number of clusters (K) in K-means clustering, for tuning the perplexity values as well. As shown through the formulation in the method section below, the Silhouette score computes the average of every point’s distance to its own cluster (cohesiveness) than to the other clusters (separateness) and is defined such that its value lies in the range of −1 to 1. A score of 1 is most desirable indicating perfectly separated clusters with clearly distinguishable features. A positive value generally indicates acceptable clustering while negative values are unacceptable for distinguishable clustering. The cohesiveness and separateness of clusters are generally measured based on Euclidean distance. Since the clusters here are identified on a reduced low dimensional t-SNE space, computing the score on this space (*S_ld_*) alone may be misleading. This is particularly true when using sub-optimal parameters that often clump the points randomly during the dimensional reduction step by t-SNE. Therefore, it is important to measure the quality of clustering with respect to the original distance in the high dimensional space (*S_hd_*), in addition to that in the low dimensional space. The integrated score (*S_ld_ ∗ S_hd_*), therefore, adds value to the estimated clustering efficiency in terms of reliability.

### B. t-SNE for clustering A *β*42 conformational ensembles

#### 1. t-SNE identifies the clustering pattern intrinsic to the A β42 ensemble

We apply our algorithm on APO and G5-bound A*β*42 all-atom MD simulations trajectories obtained from the Vendruscolo group^89^. We have used an identical set of representative frames for clustering as in the original work (35000 frames from each ensemble) where each system was simulated for 27.8 (*µ*s). Furthermore, to be consistent, we represent the conformations similarly by inter-residue Lennard-Jones contact energies. We used the distance between all pairs of conformations from the RMSD of the contact energies and feed that into our t-SNE pipeline. In the case of A*β*42 (APO and G5-bound), the calculated Silhouette score for a range of K and perplexities indicates a positive value with respect to both the distances at the low dimensional space (S*_ld_*) as well as at the high dimensional space (S*_hd_*) (Table S1 & S2) suggesting reliable clustering. This can be compared against the large negative score (−0.6) with respect to the high dimensional distance, obtained for the classical GROMOS-based clustering, which indicates that the conformations are grouped into wrong clusters. In Fig. 1(a,b), we report the integrated score (S*_hd_*\*S*_ld_*) as measured for the clusters in APO and G5-bound ensembles of A*β*42. In both cases, the Silhouette score clearly identifies an optimal cluster size (30 in the case of APO trajectory and 40 in G5 bound trajectory). The identification of clear minima in this parameter space suggests the t-SNE is able to identify a clustering pattern that is intrinsic to the underlying ensemble structure and corresponds to the true number of metastable structures. At these optimal values, we find that the low-dimensional t-SNE map shows discrete clusters in both APO and G5-bound ensemble Fig. 1(c,d) Whereas at sub-optimal values, the identified clusters either encompass different pieces together in a single cluster (for example at P=50; K=20 in APO) or break into multiple clusters of similar conformations (at P=350, K=100 in APO system).

**FIG. 1:**
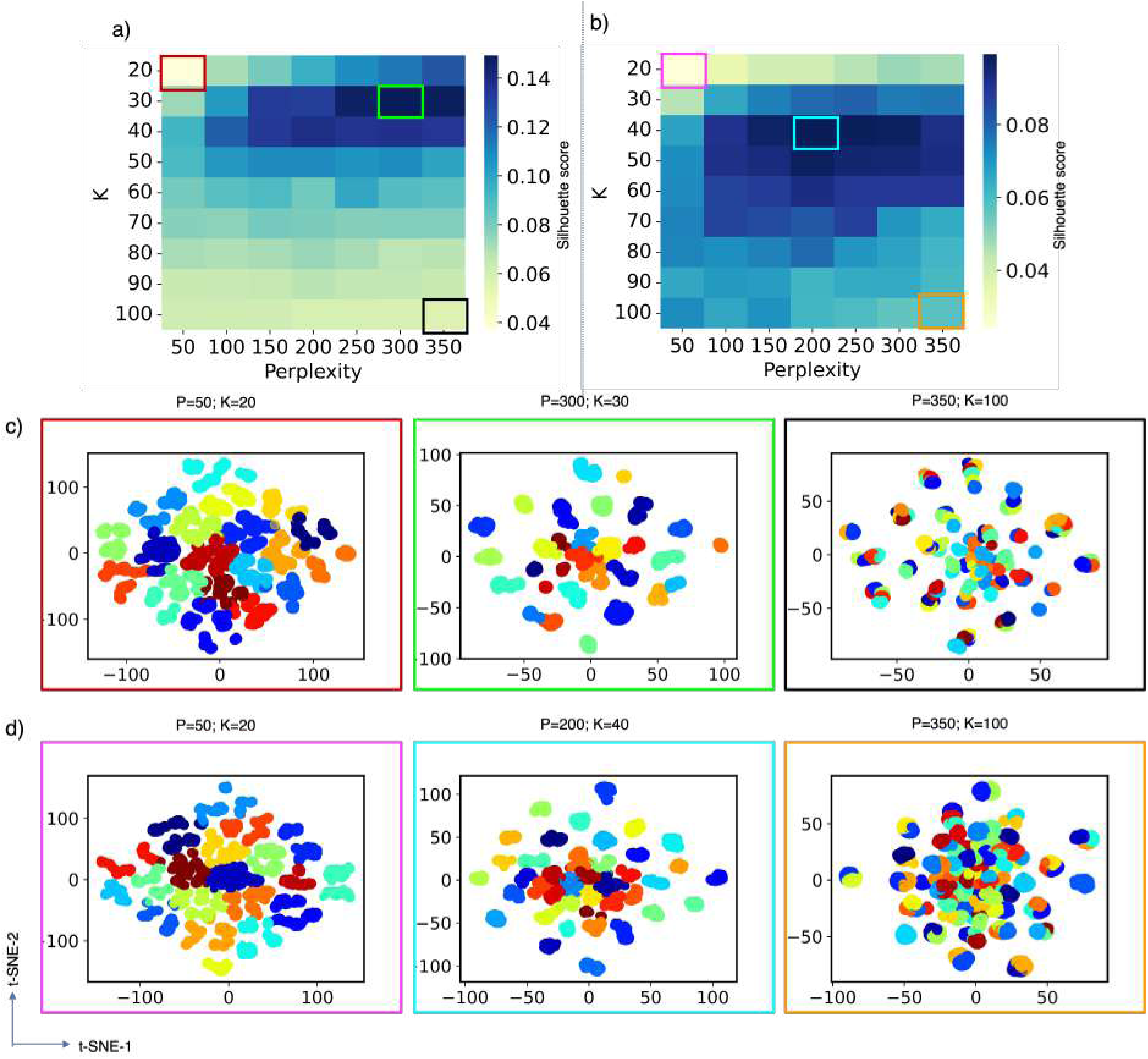
Hyperparameter optimization based on integrated Silhouette score for the (a) APO, and (b) G5-bound ensembles of A*β*42. The t-SNE maps obtained with selected optimal (green and cyan squares) and sub-optimal (Red, Black, Pink, and Orange squares) values of the perplexity and number of clusters K are shown in (c) and (d) for APO and G5 bound ensembles. The maps illustrate how these parameters affect clustering efficiency. In t-SNE projections with sub-optimal parameter values that lead to too few clusters (Red and Pink squares), we observe clearly distinguishable groups of points merged into single cluster assignments. In t-SNE projections with sub-optimal parameter values that lead to too many clusters (Black and Orange squares), we observe indistinguishable groups of points merged into different cluster assignments.

#### 2. Clustering reveals ordered sub-states within disordered A β42 ensemble

Once the optimal number of clusters for a given data set is decided using the prescriptions described above, we inspect the uniqueness and homogeneity of individual clusters by back-mapping to the conformations in the bound and unbound ensembles. Fig. 2 shows the conformations within each cluster of A*β*42 ensemble indicating unique topology and secondary structural architecture. To quantify this observation, we plotted the distance maps between conformations before and after clustering. Please see Fig. S3–S5 and subsection titled *”Estimation of homogeneity”* in SM). The results show that the clusters obtained with optimal parameters indeed yield better homogeneity than that obtained with sub-optimal parameters.

**FIG. 2:**
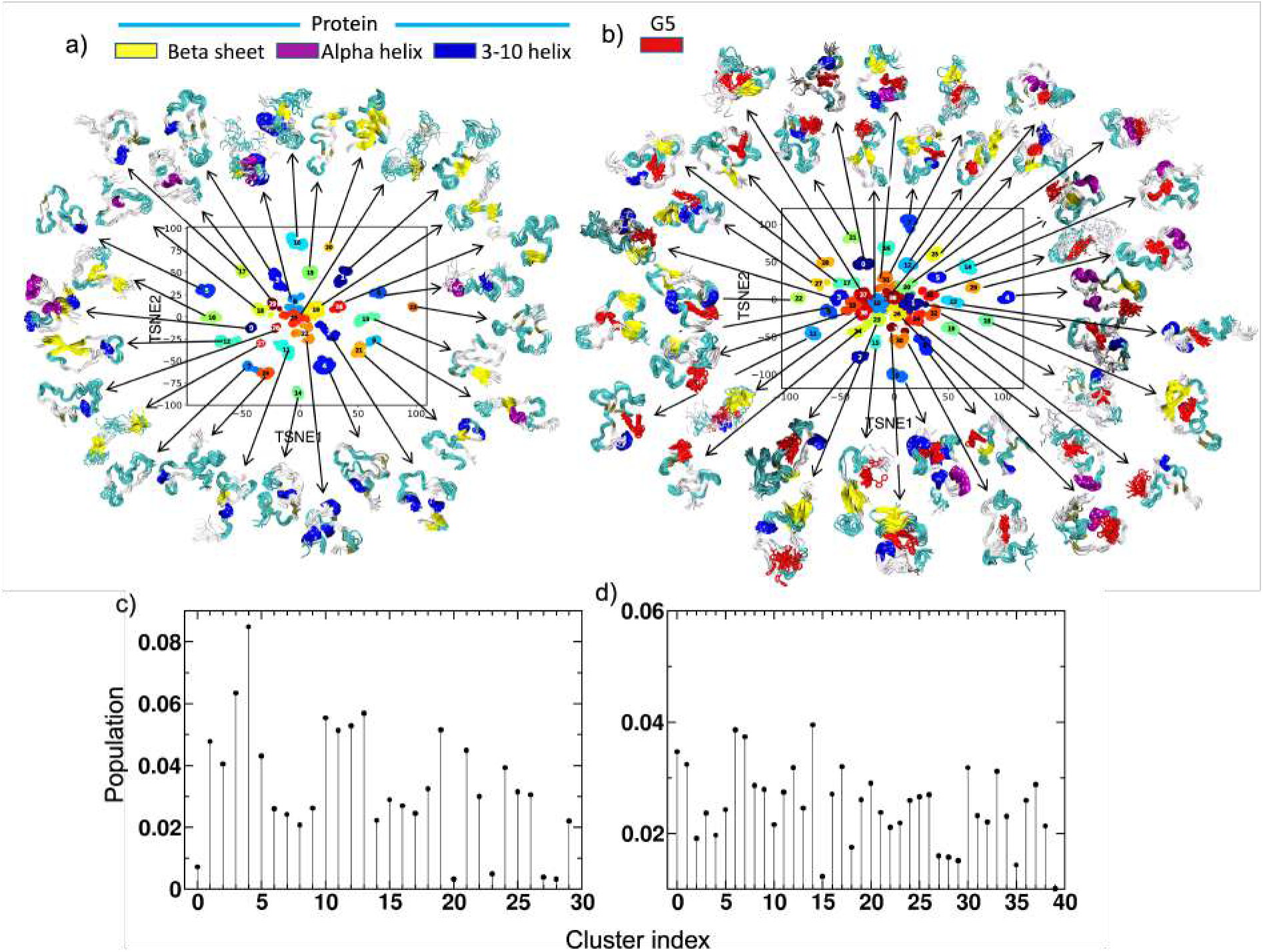
t-SNE based conformational clustering of A*β*42 ensembles in the absence and presence of G5 in a and b respectively. The cluster-wise population statistics is shown in Fig 2c and d.

More interestingly, though the G5 bound conformational ensemble was clustered only based on the similarities of protein conformations, the ligand is shown to have a specific binding orientation with the protein within each cluster (Fig. 2(b)). This result sheds light on the hidden ordered features in a disordered IDP ensemble, which can confer specificity for ligand binding. The ability of t-SNE to cluster a seemingly disordered ensemble into substates with distinct structural features and ligand binding modes suggests that one could reduce a library of tens of thousands of A*β*42 conformations to a small number of structures to screen for potential interacting ligands. This will aid in a high throughput structural and statistical analysis of IDP ensemble data and greatly aid our fundamental understanding of disorder-function relationships and in the design of therapeutic drugs for IDP molecules.

#### 3. Insights into the binding properties of Aβ42 with G5

The cluster-based population statistics of different metastable conformations have been analyzed and shown in Fig. 2(c,d). The results indicate that the distributions are more equally probable in the case of the ligand-bound ensemble than in the APO state. From this population distribution of different metastable conformations, we have estimated that the Gibbs conformational entropy (−∑(*p ln p*)) of the G5-bound ensemble is larger than the APO ensemble (Fig. S6 in SM). The number of optimal unique conformations (30 in APO versus 40 in G5-bound) and their respective Silhouette score (in high dimension space, 0.21 versus 0.15) (Table S1 and S2) also suggest consistent observation. Taken together, these results further corroborate the entropic expansion on ligand binding as deduced from the earlier studies^89^. Though the ligand has very specific binding geometry within each cluster, they vary significantly across the different clusters. We show the contact probabilities of G5 with individual protein residues in Fig. 3(a) for individual clusters. We also plot the residue-wise contact probabilities using the total trajectory, which provides averages without clusters (Fig. S7 in SM). As indicated by the figures, the G5 preferentially binds to aromatic residues such as Tyr/Phe (residue numbers 10, 19, 20) and hydrophobic residues such as Ile/Val/Met (residue numbers 31, 32, 35, 36). The interactions of G5 with these aromatic and hydrophobic residues potentially disrupt tertiary contacts between these residues in the A*β*42 ensemble, thus limiting the stabilization of transiently ordered A*β*42 conformations and increasing the heterogeneity and conformational entropy of the ensemble. To further quantify how the contacts of G5 at diverse locations affect the interaction strength, we applied a high throughput numerical technique called molecular mechanics with generalized Born and surface area solvation (MM/GBSA) to estimate the free energy of the binding of ligands to proteins^93,94^. Our MMGBSA-derived binding scores are shown in Fig. 3(b). We see that the G5 binds at relatively equal strength in multiple clusters. But interestingly, we also noted a few of the clusters (cluster numbers 14, 29, and 30) that show statistically stronger binding than the others. More interestingly, these same clusters consist of a relatively larger population in the ensemble than the other conformers. The protein residues involved in binding in these selected clusters along with their energy contributions to the total energy as plotted in Fig. 3(c) and the conformational binding-geometry for the cluster that exhibits the most favorable MM/GBSA binding is shown in Fig. 3(d,e). In Fig. S8 in SM, we also show the same data (binding geometries and residue-wise interactions) for the two other clusters, which show the second and the third-best MM/GBSA scores. Our analyses reveal that ligands interact with multiple favorable sites simultaneously, which indicates that even a partially collapsed or ordered state of an IDP can provide a specific binding pocket for small molecule interactions. These unique insights gained as a result of high-fidelity clustering can be leveraged for future IDP-drug designing with conventional strategies utilized to target ordered binding sites in folded proteins.

**FIG. 3:**
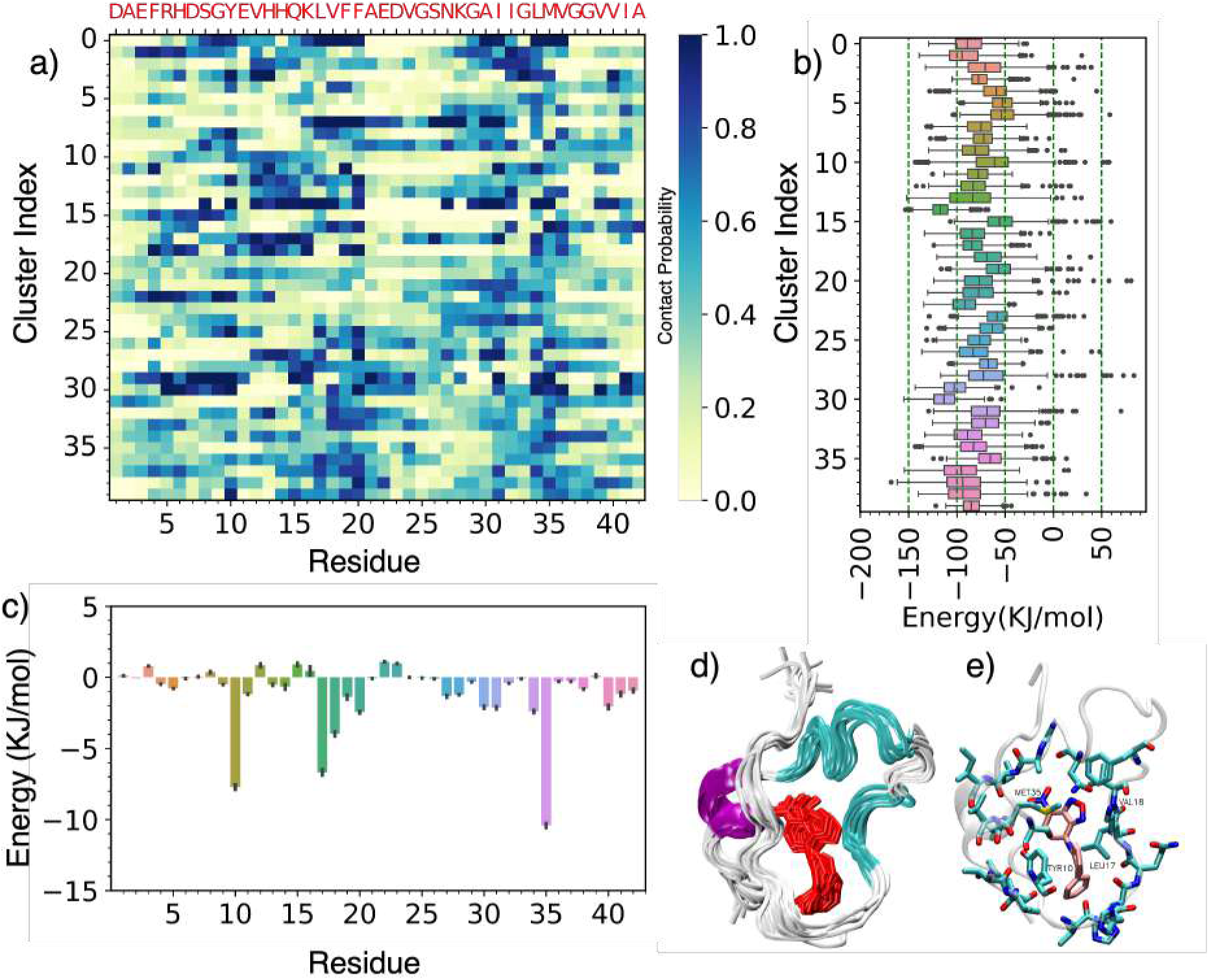
Cluster-wise inter molecular contact probabilities and their respective binding energy as measured using MMGBSA analysis are shown in (a) and (b) respectively. For the cluster that shows the most favorable binding (cluster no: 14)), we have shown the residue-wise decomposed energy contribution in (c) with error bars representing 99% confidence interval of the estimated mean. The superposition of ten central conformations from this specific cluster is shown in (d) and the interacting residues are shown in stick representation in (e)

### C. t-SNE for clustering *α*-synuclein conformational ensembles

#### 1. t-SNE reveals distinct conformation sub-states despite extreme structural plasticity

Next, we apply our clustering algorithm to characterize the conformational ensemble of the prototypical IDP *α*-synulcein (*α*S). *α*S is a longer IDP than A*β*42, consisting of 140 amino acids, and has a substantially less ordered, or more ”fuzzy”, conformational landscape with virtually no experimentally detectable residual secondary structure propensity. We also apply our t-SNE clustering algorithm to cluster the conformations of a C-terminal fragment of *α*S, containing residues 121-140, which we refer to as ”*α*S C-term”. These residues were shown to have the highest affinity to a family of small molecule ligands based on the structure of the Rho protein kinase inhibitor Fasudil by both NMR experiments and unbiased MD simulations of a full-length *α*S construct^91^. The t-SNE projection of full length *α*S produces a crowded map, with only a few segregated clusters of points visible(Fig. S9a). In the case of *α*S C-term, t-SNE projections produce a single continuous grouping, or ”blob”, of points, with no clearly distinguishable subsets of data points regardless of the perplexity value used, suggesting extreme heterogeneity and almost no detectable order in its conformational landscape (Fig. S9b). This distribution of points in the low dimensional t-SNE projection suggests *α*S C-term may be described by a broad and relatively flat energy surface with very few barriers or local minima. This is in stark contrast to the substantially more discernible t-SNE projections data of A*β*42 seen in Fig. 1.

In order to obtain a better sense of the conformational diversity of *α*S and *α*S C-term, we examined the pairwise RMSD between conformations in both ensembles in Fig S10. Here, we observe that the conformational states rapidly exchange among themselves, which in turn creates a very cluttered distance map of the original trajectory. This is shown in the first subplot for full *α*S in Fig. S10(a) and for the C-terminal (C-term) peptide in Fig. S10(b). This suggests there are very few intrinsic groupings of these conformations in the high dimensional space, which is consistent with the t-SNE projections seen in Fig. S9. When we apply our t-SNE clustering approach and scan values of perplexity and cluster size, we observe substantially worse Silhouette scores relative to those obtained for A*β*42, with values very close to 0, indicating poor clusterability of these ensembles. However, we find that in this relatively continuous distribution of conformations, we still observe some positive Silhouette scores, though with very small magnitudes, suggesting some limited success in projecting onto a lower dimensional manifold. A small magnitude positive Silhouette score can indicate that most data points are on or very close to the decision boundaries between neighboring clusters. In such cases, scanning values of Silhouette scores as a function of perplexity values and the number of clusters may not locate a clear maximum in this parameter space (Fig S11). In the case of full-length *α*S we observe that the Silhouette score continues to increase with a number of clusters beyond an undesirably large number of clusters (over 100) that becomes difficult to structurally interpret. In the case of *α*S C-term, where we see almost no separation of points in lower dimensional t-SNE projection, we observe that the Silhouette score is at a maximum with two clusters, and decays to zero as the number of clusters increases beyond 2. We, therefore, observe that our procedure for scanning the parameter space of perplexity and cluster size is less successful for the substantially more continuous distribution of conformations observed in simulations of *α*S and *α*S C-term.

Nevertheless, we attempted to determine if t-SNE projections of the conformational ensembles of *α*S and *α*S C-term onto a lower dimensional space can provide interpretable structural insights into these ensembles. Due to the nature of the projection data, we do not use the usual for an optimal Silhouette score. Instead, we focused on a tractable number of clusters and manually choose the perplexity and number of clusters in an effort to achieve a reasonable degree of structural homogeneity within cluster assignments. To assess the interpretability of t-SNE projections with low Silhouette scores, we have examined the structural properties of clusters generated with K=50 and perplexity=400 for full-length *α*S and K=20 and perplexity=1800 for *α*S C-term (Fig. 4, Fig. S11). The clusters produced with these values effectively divide the continuous distribution of points in the t-SNE projection space into contiguous regions with no clear separations in the lower dimensional projection. We then inspect the conformational homogeneity of the structures in each cluster to determine if this discretization provides interpretable structural insights. Despite the lower Silhouette scores, we observe substantial conformational homogeneity within these cluster assignments as assessed by the visualization of the conformational states (Fig 4) and pairwise RMSD between clusters (Fig S10). This suggests that our t-SNE low dimensional projection preserves local structural properties of IDPs well even when distinct clusters of data points are not apparent based on the low dimensional t-SNE projections and Silhouette scores.

**FIG. 4:**
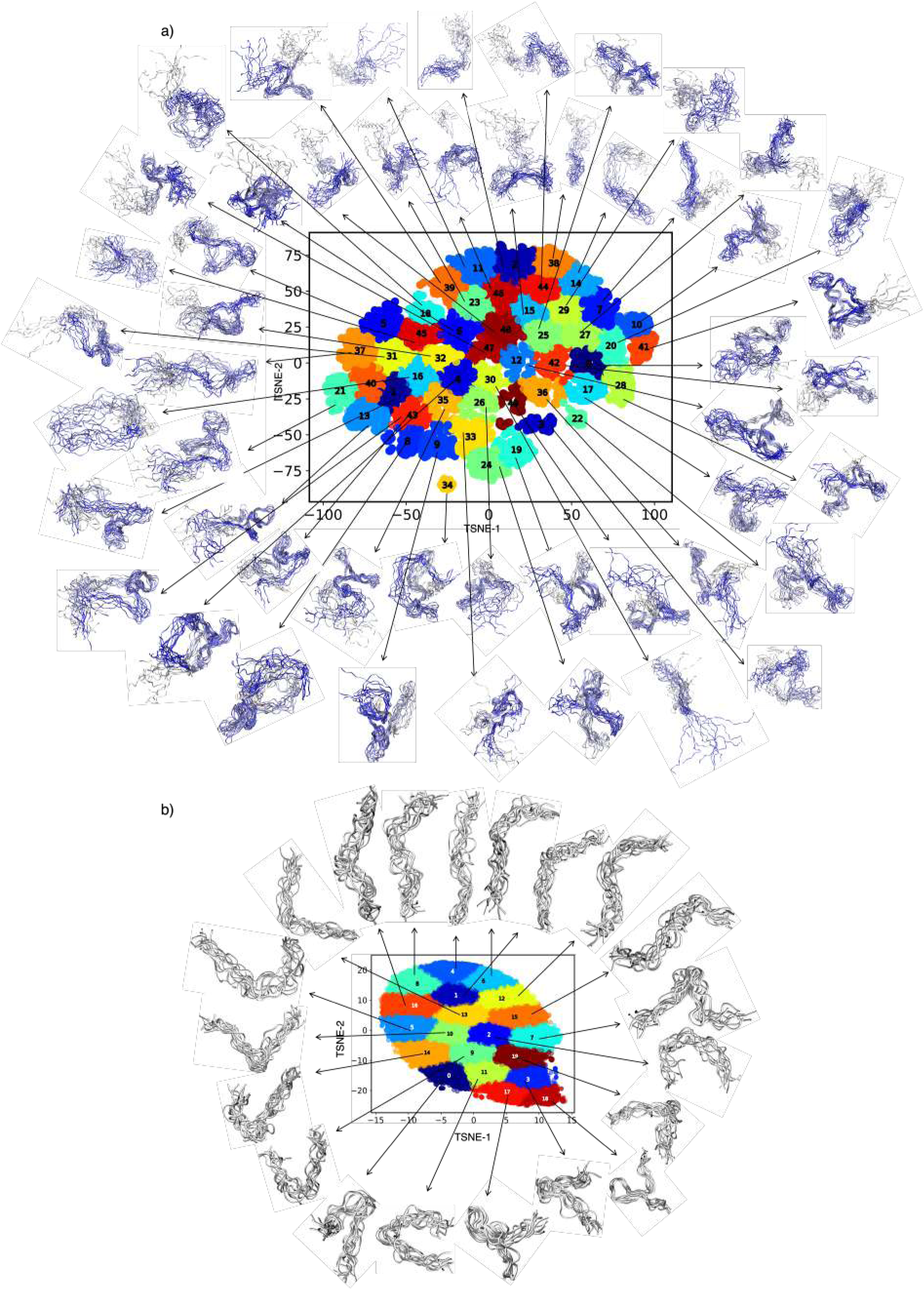
t-SNE based conformational clustering of (a) full-length α-synuclein (140 residues) and (b) a 20 residue C-terminal fragment of a-synuclein.

Visual representations of the conformations in the 50 clusters of full-length *α*S system and 20 clusters of apo C-term *α*S system are shown in Fig. 4(a,b). In spite of the extreme heterogeneity of the conformational space of the *α*S and *α*S C-term ensembles, and relatively continuous distribution of points in the low dimensional t-SNE projections, we find that our clustering method clearly partitions *α*S and *α*S C-term conformations into clusters with unique and relatively homogeneous conformations. While the conformations do not contain any secondary structure and do not collapse to form rigid pockets as in the case of the A*β*42, we still observe substantial order within each of the clusters. Interestingly, we find that the conformations of C-term *α*S peptide span a range of conformational states that vary from fully-extended rod-like shapes to acutely bent hairpin-like conformations (4(b)) and presents all intermediate bending angles between these two extremes. To illustrate this feature, we have presented the clusters in a sequence, arranged based on the average bend angle measured between the C*α* atoms of residues 121, 131, and 140 (Fig. 5(a-c)), which make up the C-terminal, middle and N-terminal residues of the peptide, respectively. Henceforth, we will refer to this simply as the ”bend angle”. We have plotted the distribution of the bend angles observed in each cluster in Fig. 5(c). An interesting and valuable by-product of this high-fidelity clustering is that it seems to inform a single collective variable that uniquely defines the various conformations across clusters. This collective variable may be useful for running computationally efficient biased simulations of this system.

**FIG. 5:**
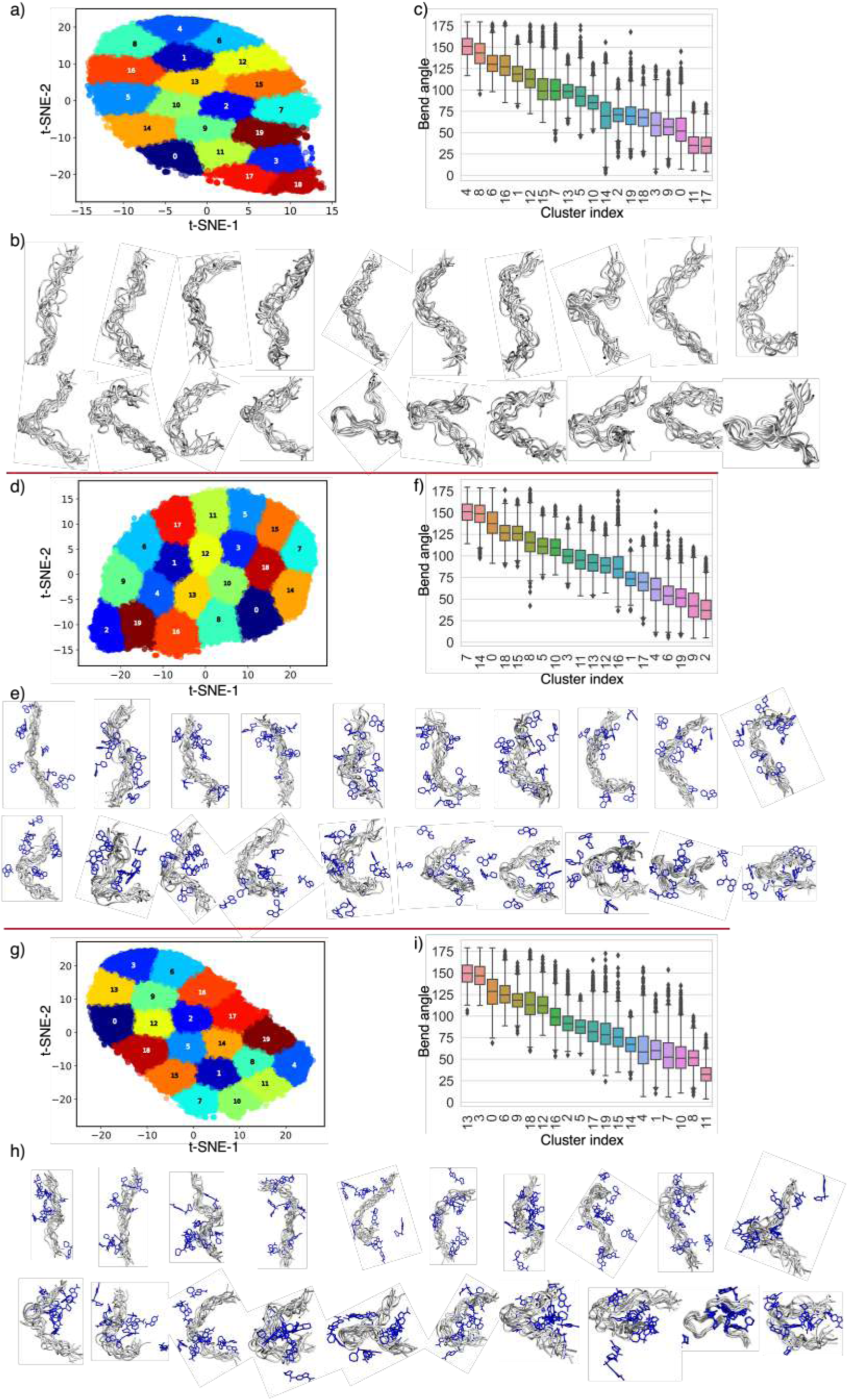
t-SNE based conformational clustering of APO, fasudil-bound and ligand47-bound ensembles of a C-terminal fragment of α-synuclein are shown in Top, middle and bottom respectively. The conformational subspace of the t-SNE projections is subdivided into 20 clusters (Fig 5a, 5d and 5g). The structure of these conformations within each cluster shows a relatively homogenous distribution of structures (b,e and h). The clusters of conformations are arranged based on the bend angle calculated on the *Ca* atoms of residues 121, 131, and 140 and displayed in order of a decreasing bend angle. The distribution of the bend angle in each cluster is shown as a box plot (in c,f, and i) for the apo, Fasudil-bound, and Ligand47-bound ensembles.

#### 2. Characterization of ligand bound ensembles of the C-terminal αS peptide

We next used our t-SNE clustering approach to quantify the effects of small molecule ligand binding on the conformational ensemble of *α*S C-term. We have chosen to analyze the effects of binding two ligands, the small molecule Fasudil and a previously identified higher affinity *α*S ligand (ligand 47), on the conformational ensemble of *α*S C-term. We first generated t-SNE maps for all conformations of *α*S C-term in the presence of fasudil or ligand 47 for different values of perplexity as shown in Fig. S12. Similar to the low dimensional projection of conformations observed in the APO simulation of *α*S C-term, we observed that the t-SNE projections of both ligand-bound ensembles produce a continuous distribution of points with no clearly distinct subsets of points. We then clustered the conformations using the same number of clusters K=20, and selected perplexity values for each ligand-bound ensemble that achieved the maximal silhouette score for K=20 (perplexity=1200 and perplexity=1100 for the fasudil bound ensemble and ligand 47 bound ensemble respectively) (Figure 5d-e and 5g-h). As was the case with the APO ensemble, we find that these clusters partition conformations of *α*S C-term by the previously defined bend angle between the C*α* atoms of residues 121, 131, and 140 (Figure 5f and i).

Unlike the localized binding of G5 with individual metastable states of A*β*42, the binding of fasudil and ligand 47 is not localized to a specific region of the peptide within each cluster (Fig. 5(e,h). We quantified the inter-molecular contacts between fasudil and ligand 47 with *α*S C-term in figure 6a,b. We also quantified the fraction of specific interactions in each cluster where a protein residue forms a hydrophobic contact, an aromatic stacking interaction, a charge-charge contact, or a hydrogen bonding interaction with the ligand utilizing reported geometric criteria (Figure S13 and S14)^91^.

**FIG. 6:**
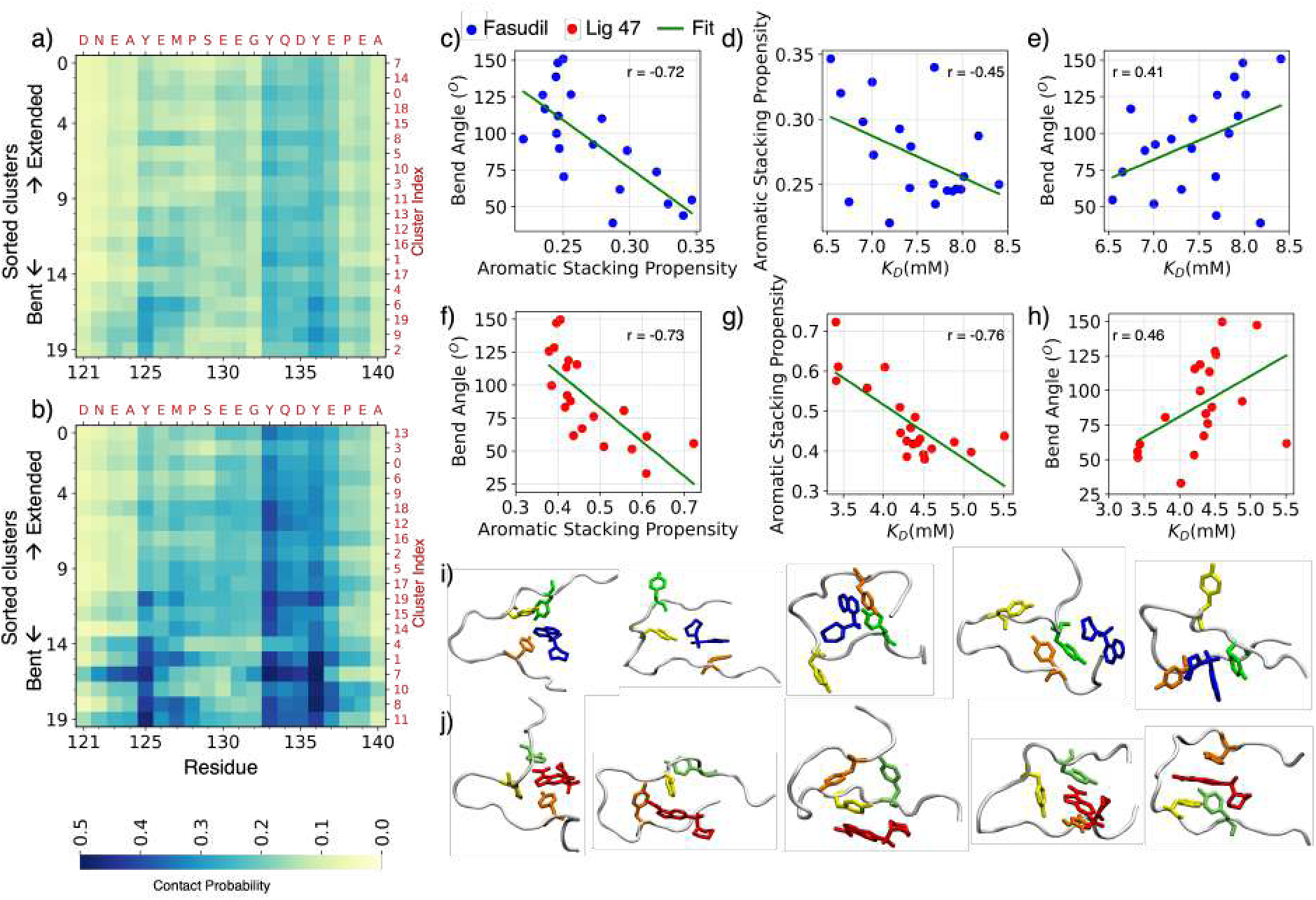
Per-residue intermolecular contact probabilities between *α*S*_C−term_* and fasudil and *α*S*_C−term_* and ligand 47 observed in each cluster are shown in (a) and (b) respectively. The clusters are sorted in the decreasing order of bend angle and the actual cluster indices are indicated in the alternate Y-axis in red. Figures 6c-h represent the correlations among the average bend angle, total aromatic stacking propensity, and dissociation constant, (K*_D_*), measured from individual clusters. The corresponding Pearson correlation coefficient is indicated within each plot. Representative snapshots from the top 5 clusters containing acutely bent hairpin-like conformations of Ligand bound αS*_C−term_* illustrating how the bent conformations orient the aromatic side chains of Tyr-125, Tyr-133, and Tyr-136 towards better stacking interaction with Fasudil (i) and Ligand 47 (j) that in turn lead to better inter-molecular affinity. The snapshots from left to right were taken from cluster numbers 4, 6, 19, 9, and 2 in the case of Fasudil-bound *α*S*_C−term_* (i) and cluster numbers 1, 7, 10, 8, and 11 in case of Ligand-bound αS*_C−term_* (j).

For both ligands, we observe that the maximum contact probability with any residue is only 0.5. We note that the contact analyses are carried out with the full trajectory of bound+unbound frames, which could be the factor for lower values. Interestingly, we observe the same trend even when we analyzed only the bound frames of the *α*S C-term trajectories in all the clusters (Fig. S15). Further, we observe relatively similar intermolecular interaction profiles across clusters, with relatively smaller deviations, and the contacts are primarily centered around the three aromatic residues of *α*S C-term (Y125, Y133, and Y136) illustrating that the same sets of intermolecular interactions are accessible regardless of the distribution of bend angles in each cluster. We do not observe any specific sets of intermolecular interactions, such as specific charge-charge contacts or aromatic stacking interactions, that are only present in a subset of clusters. These relatively lesser contacts at any specific residue and similar interaction profiles across clusters are consistent with the previously proposed ”dynamic shuttling” mechanism of IDP small-molecule binding, where small molecule ligands transition among a heterogenous ensemble of binding modes based on the geometric proximity potential sidechain and backbone pharmacophores^91^.

Examining the intermolecular interaction profiles of the two ligands, we observe that ligand 47 appears to have substantially higher fractions of aromatic charge contacts (Fig. S13 vs Fig. S14) than Fasudil. Surprisingly, the population of aromatic stacking interactions seems to be dependent on the bend angle of *α*S C-term in both the ligands. The clusters with acutely bent conformations mostly have higher aromatic stacking propensity. The Pearson correlation between the cluster-wise average bend angle and total aromatic stacking is very high (−0.7), as shown in Fig. 6c and f. To us, this was a very unique and non-obvious observation that manifested itself due to our clustering exercise. To obtain more detailed insight into the binding modes of Fasudil and ligand 47 we examined the relative affinity of the ligands to each cluster. Considering both apo and bound frames in clustering enabled us to calculate the fraction of bound frames in each cluster, and report simulated K*_D_* values for each cluster as reported previously^91^ (Figure S16). The K*_D_* values of Fasudil and ligand 47 range from 6.5mM-8.5mM and 3.5-5.5mM across clusters, respectively. Though there are only small deviations in the K*_D_* values across the clusters, we notice that the K*_D_* is significantly lesser in clusters with acutely bent conformations and high aromatic stacking propensity (Fig 6d,e,g,h). The strong correlation between these values suggests that the bent conformations provide substantially more compatibility toward binding by orienting the aromatic residues (Y125, Y133, and Y136). Representative snapshots from the top 5 bent clusters of Fasudil and Ligand 47 bound *α*S C-term are shown in Fig. 6i and j. This is a very exciting result to us since it provides a relationship between the relative curvature of the *α*S C-term backbone and the accessibility of specific intermolecular interactions. This relationship is much stronger in the higher affinity ligand 47, suggesting that exploiting a coupling between conformational substates and the accessibility of specific intermolecular interactions such as aromatic stacking may be useful for designing higher affinity ligands for disordered IDP ensembles.

Since the conformational ensembles of *α*S C-term were obtained from unbiased MD simulations, we can assess the kinetic stability of the conformations in the reported clusters by calculating the transition probabilities between clusters at different lag times (Fig. S17). Here we observe that most clusters in the APO *α*S C-term are not well defined in terms of kinetic stability. Even at these short timescales, for bend angles greater than 70*^o^* there is little memory of cluster assignment in the trajectory, and no noticeable pattern of transition probabilities between clusters. This pattern of transition probabilities is consistent with the notion of a broad and flat free energy surface with few local minima. We notice that there seems to be elevated kinetic stability for *α*S C-term conformations with small bend angles (¡70*^o^*) at short lag times. This suggests a slightly more rugged conformational free energy surface for hairpin-like conformations, which are likely stabilized by sidechain interactions between residues more distant in sequence. We observe however that the kinetic stability of hairpin-like conformations of apo *α*S C-term is not observed at longer timescales, suggesting that the local free energy minima of hairpin conformations are fairly shallow. We observe a similar pattern of kinetic stability of *α*S C-term clusters observed in the presence of fasudil and ligand 47.

Lastly, we compare the shift in the populations of conformational states of *α*S C-term and A*β*42 in the presence and absence of small molecule ligands by projecting the conformational ensembles of apo simulations and simulations in the presence of ligands onto a single t-SNE projection for each protein (Fig S18). In the case of *α*S C-term, we observe that the ligand-bound and apo ensembles are nearly indistinguishable in the lower dimensional t-SNE projection. This is in severe contrast to the behavior exhibited by A*β*42 APO and ligand-bound t-SNE projections as shown in Fig. S18 (b). The map in Fig. S18 (b) clearly shows that the APO and bound A*β*42 ensembles have clusters that are distinct with only a few regions showing overlapping projections.

### D. Scope and limitations of t-SNE method with IDP-clustering

Unlike the commonly used projection techniques such as PCA and MDS, t-SNE optimization is non-convex in nature with random initialization that produces different sub-optimal visual representations at different runs. While the physical interpretation of t-SNE projections seems daunting, this affects mainly the global geometry and hierarchical positioning of the clusters and not the local clustering pattern. We illustrate the consistency in local clustering upon different runs with different random initialization by quantifying the Silhouette score and mutual information of clusters in Table S4. Moreover, finding a single optimal global geometry of the IDP dataset is often not possible owing to their extreme heterogeneity with almost equal transition probability between different clusters. However, if one necessitates the global preservation, tuning the perplexity^95^, and other parameters like Early exaggeration and Learning rate, initializing with PCA and Multi-scale similarities could be helpful^75,96^. In addition, some of the variations of t-SNE methods such as h-SNE can also be helpful^97^.

Another factor that should be considered while using t-SNE on ultra-large datasets is the associated computational cost. Analyzing large data sets with t-SNE (beyond n *≫* 10^6^) is not only computationally expensive (scales with *O*(*n*^2^)), but also suffers from slow convergence and fragmented clusters. If the computational cost becomes formidable, one could use methods such as Barnes-Hut approximation^98^ and the FIT-SNE method to accelerate the computation. In short, Barnes-Hut approximation considers a subset of nearest neighbors for modeling the attractive forces, and the FIT-SNE method relies on a fast Fourier transformation, which reduces the computational complexity to *O*(*NlogN*) and *O*(*N*), respectively. To mitigate the slow convergence and fragmentation of clusters, it is often desirable to run t-SNE on a sub-sample of the trajectory that includes all unique populations and then projects the rest of the points onto the existing map.

In line with discussing the possible pitfalls of the t-SNE method, it is also understood that adding a new data point onto the existing t-SNE map can lead to erroneous interpretation as the method is essentially non-parametric and does not directly construct any mapping function between the high dimensional and low-dimensional space. Recent extensions of the method in combination with deep neural networks allow for parametric mapping^75,99^ and could be tried if such a situation can not be avoided. Moreover, the possibility of out-of-sample mapping with parametric t-SNE can be explored further for driving simulations from one state to another and to match experimentally known values. For instance, in such cases, the similarities can be obtained from NMR chemical shifts or from SAXS intensities. From that perspective, t-SNE as an integrative modeling tool looks very promising.

## III. Conclusion

In spite of the well-established knowledge of the inherent conformational heterogeneity in an IDP ensemble and despite advances made in accurately determining the ensemble conformations using integrative approaches, successful application of IDPs to drug targeting is limited. The main reason behind this is the lack of accurate classifications of the conformational ensemble. Our algorithm provides that tool where several thousands of structures can be grouped into representative sets of the distinct and tractable conformational library, with unprecedented quality and performance. We introduced new metrics for choosing optimal hyper-parameters of the algorithm and for validating the homogeneity in the resultant clusters. The accessibility and generality of the framework enable faithful clustering for broader applications without requiring expert domain knowledge of the underlying data.

We demonstrated the approach on a*β*42 and *α*S ensembles under free and ligand-bound contexts. Our results provide important insights into the ordered meta-stable structures present in the IDP ensembles and their binding mechanism to small molecule ligands. The two IDPs studied here exhibits vastly different mechanism of small molecule recognition: while the Abeta42 has distinct binding pockets in different metastable structures that bind uniquely with the G5 molecule, the binding of Fasudil and ligand 47 with *α*S C-term follows the ”dynamic shuttling” within all the metastable states. Yet the residue preference across the two ligands with *α*S and the G5 with the A*β*-42 is strikingly similar for the aromatic residues. This was also observed in another recent study for small molecule binding in p27^100^. Designing ligands that target these residues could be a common strategy for IDPs. The results of t-SNE based clustering exercise on *α*-S reveal one of the most interesting and non-obvious learning about the emergence of the possible role of peptide local curvatures, besides the weak chemical specificity, as sites of ligand binding. Our tool also makes it very convenient to generate sub-groups of similar conformations for long IDPs, whose full conformational ensemble is highly intractable for structural biophysical analyses. For example, we applied our algorithm to study how the long disordered regions of FUS protein interact with RNA molecules^101^ and this t-SNE tool allowed us to illustrate the complex RNA binding behavior of the long disordered FUS RGG repeats in an interpretable manner. Taken together, these learnings will invariably aid in carrying out *in silico* functional and drug screening studies in a rational manner, a critical next step for curing many incurable IDP-induced diseases. Identification of functionally and pathologically relevant substructure of an IDP would also open ways for reverse engineering of IDPs with functions useful in biotechnology and medicine.

## IV. Materials and Method

### A. Input for t-SNE analysis

The systems details about the trajectories of alanine-dipeptide, A*β*42, and *α*S ensembles are reported in Table 1. The conformations of the trajectories were represented by backbone dihedral angle, inter-residue LJ-interaction potential, and atomic coordinates of heavy atoms for alanine-dipeptide, A*β*, and *α*S ensembles, respectively.

#### 1. t-SNE based dimensional reduction

Given a number of observations (conformations) *n* and with *d* dimensional input features in the original space defined as *X* = *{x*_1_, *x*_2_, …, >*x_n_} ∈* R*^d^*, t-SNE maps a smaller *s* dimensional embedding of the data that we denote here by *Y* = *{y*_1_, *y*_2_, …, *y_n_} ∈* R*^s^*. Here *s ≪ d* and typically s = 2 or 3. This projection is based on the similarity and dissimilarity between conformations. The similarity or dissimilarity between the conformations in the high dimensional space is computed based on Euclidean or RMS distances. t-SNE aims to preserve the local neighborhood such that the points that are close together in the original space remain closer in the embedded space. In the original space, the likelihood of a point *x_j_* to be the neighborhood of *x_i_* instead of every other point *x_k_* is modeled as a conditional probability *p*_(_*_j|i_*_)_ assuming the Gaussian distribution centered at point *x_i_* with a standard deviation of *σ_i_*.

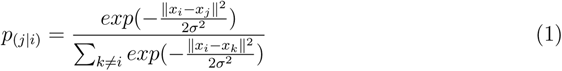

Similarly, the conditional probability in the embedded space (*q*_(_*_j|i_*_)_), with the same n points initialized randomly, is computed but now based on a t-distribution. Having a longer tail than Gaussian, the t-distribution moves dissimilar points farther away to ensure less crowding in the reduced space.

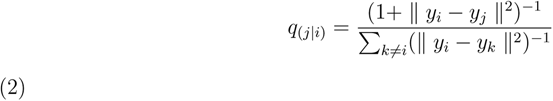

To ensure symmetry in the pairwise similarities, the joint probability is calculated from the conditional probability as follows:

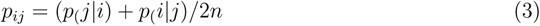

Finally, the difference between the two probability distributions, calculated as Kullback-Leibler (KL) divergence is then minimized by iteratively rearranging the points in the low dimensional space using gradient descent optimization.

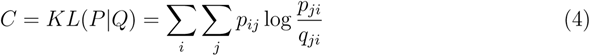

where P and Q are the joint probability distributions in the high and low dimensional space over all the data points.

The major tunable hyperparameters in t-SNE are the perplexity, learning rate and the number of iterations. The perplexity value, *P* defines the Gaussian width, *σ_i_*, in Equation 1 above such that, *log_2_P* = *H*(*P_i_*) = −∑*_j_ p_j|i_* log_2_ *p_j|i_* for all *i*. Loosely, this parameter controls the number of nearest neighbors each point is attracted to and therefore balances the preservation of similarities at a local versus global scale. Typically, low perplexity values tend to preserve the finer local scale and high perplexity values project a global view. To optimize perplexity, we ran the algorithm with varying values of perplexities and chose the one that yields a high silhouette score. The other two parameters such as the learning rate and the number of iterations control the gradient descent optimization. While we chose the default value of 200 for the learning rate, the number of iterations was chosen to be 3500, which is large enough for avoiding random fragmentation of clusters as suggested in the literature.^96^

### B. Kmeans clustering of data on the reduced space obtained from t-SNE

Kmeans clustering is the simplest unsupervised clustering algorithm that partitions the data into non-overlapping clusters. The algorithm starts by grouping data points randomly into K clusters, as specified by the user. Then it iterates through computing the cluster centroids and reassigning data points to the nearest cluster centroid until no improvements are possible. The parameter, K, is optimized by running at various values and chosen based on the maximized clustering efficiency.

### C. Optimizating the hyperparameters (*Perp* in t-SNE and *K* in k-means) using Silhouette score

Silhouette score for a datapoint i is measured by,

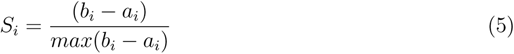

where *a_i_* is the intra-cluster distance defined as the average distance to all other points in the cluster to which it belongs. *b_i_* represents the inter-cluster distance measured as the average distance to the closest cluster of datapoint *i* except for that it’s a part of. Typically the Silhouette score ranges between 1 and −1, where a high value indicates good clustering, and values closer to 0 indicate poor clustering. A negative value indicates the clustering configuration is wrong/inappropriate.

The distance between points is usually measured in terms of the Euclidean distance metric. Since the clusters, in our case are identified in a reduced representation with t-SNE, computing the score based only on the distances in the reduced space (*S_l_d*) may be misleading, if the points are wrongly put together during the dimensional reduction step by t-SNE. Therefore, it is important to measure the goodness of clustering with respect to the original distance in the high dimensional space (*S_h_d*), in addition to that in the low dimensional space. The integrated score (*S_l_d ∗ S_h_d*), therefore, adds value to the estimated clustering efficiency in terms of reliability.

### D. Cluster-wise conformational analysis and visualization

The conformations corresponding to each cluster are extracted using Gromacs based on the cluster indices. All the conformations were used for estimating the contact probability, binding energy, and homogeneity within individual clusters. Whereas, for visualization purposes, we extracted ten representative conformations from each cluster that is closest to the corresponding cluster centroid (as identified using KD-tree based nearest neighbor search algorithm). The conformations are rendered using VMD.

Our current implementation of the model is available on the GitHub repository: https://github.com/codesrivastavalab/tSNE-IDPClustering.

## V. Author Contributions

R.A. and A.S. conceived and designed the research; A.R. performed the calculations with help from J.K; R.A., J.K., M.B., P.R., and A.S. analyzed data; R.A., P.R., and A.S. wrote the paper together with inputs from J.K., M.B.

## VI. Acknowledgments

A.R. thanks the Wellcome Trust DBT India Alliance for Early Career Fellowship (Grant number: IA/E/18/1/504308). A.S. thanks the Department of Science and Technology (DST) of India for the early career grant (SERB-ECR/2016/001702). A.S. also thanks the DST for the National Supercomputing Mission grant (DST/NSM/R&D HPC Applications/2021/03.10). Computational support from the high-performance computing facility ”Beagle” setup from grants by a partnership between the Department of Biotechnology of India and the Indian Institute of Science (IISc-DBT partnership program) is greatly acknowledged. AR and AS are grateful to the SciNet HPC Consortium, ComputeCanada for their generous computational support. P.R. and J.K. are supported by the National Institutes of Health under award R35GM142750.

## Supplemental Material

**FIG. S1:**
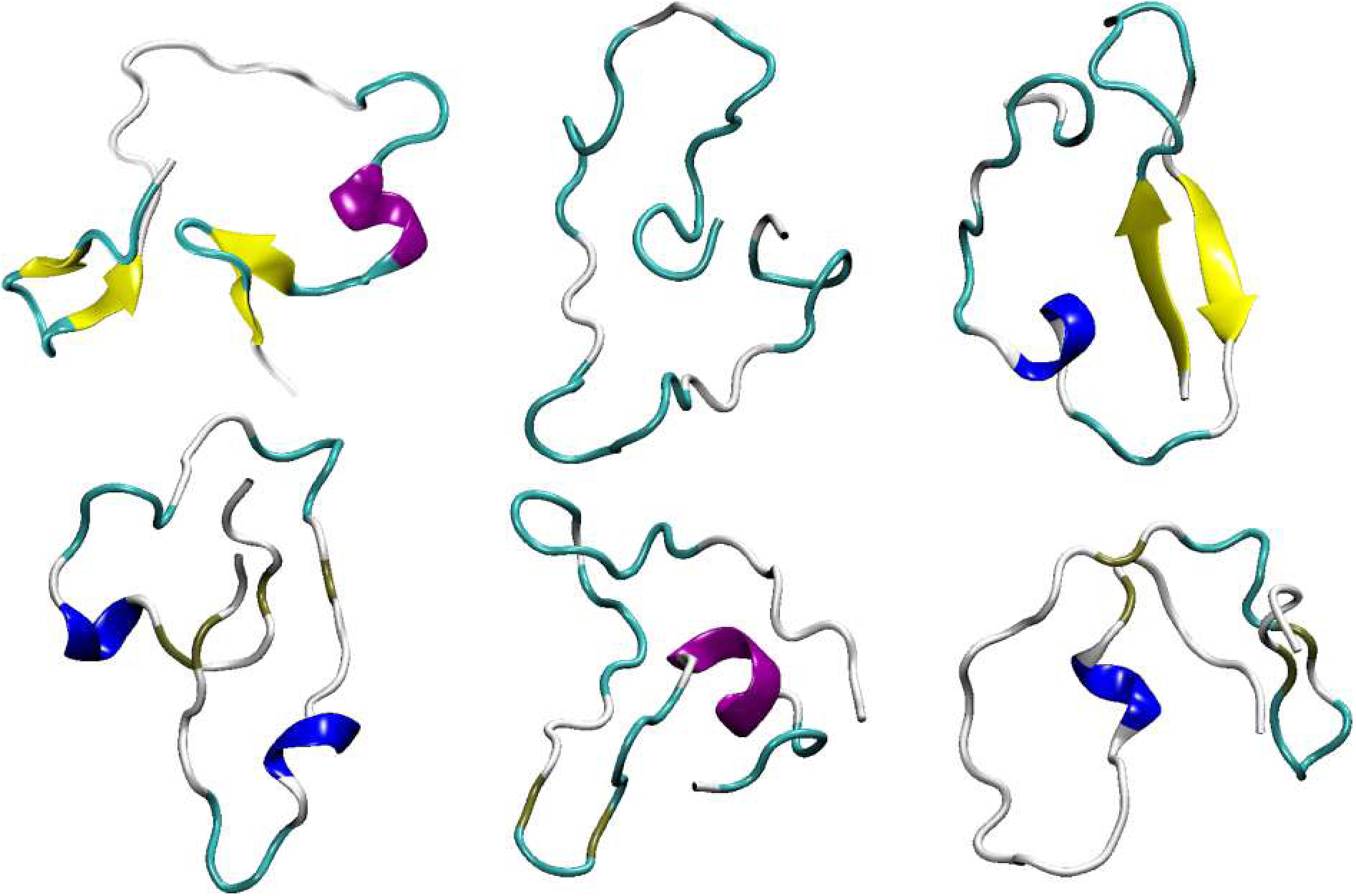
Snapshots of several conformations of A*β*42 with the same Rg values (1.10 nm) but have different structures altogether. It is evident from this illustration how the use of such low-dimension collective variables (CVs) could lead to ambiguous classification.

### A Physical intuition into t-SNE-based clustering algorithm using alanine dipeptide as a model system

We first employ the t-SNE method on the alanine dipeptide (ADP) trajectory where we compare the results with the well-known 2D Ramachandran plot using dihedral distance as dissimilarity score. Ramachandran plot is also called *ϕ − ψmap* due to the backbone dihedral angles along the peptide bond^1,2^. Here, we can fix the number of clusters to four (K = 4) based on the four known sub-regions of the Ramachandran plot namely beta-sheet, PPII, right-handed *α* helix, and left-handed *α* helix (Fig. S2(a)). The left-handed *α* helix region lies separately in the second half of the *ϕ* dihedral axis whereas the beta-sheet and the right-handed *α* helical regions occupy the first half of the *ϕ* dihedral axis. To quantify the goodness of clustering, we calculate the Silhouette score^3^ on the raw data and arrive at a score of 0.55 for the 2D map. Of note, when the numbers of clusters are not known a priory unlike the ADP system, we have a prescription that makes use of silhouette score with the t-SNE perplexity values to find the optimum number of clusters.

The second feature of t-SNE is a tuneable parameter called “perplexity,” which (loosely) dictates how to balance attention between local and global aspects of your data. The parameter is, in a sense, a guess about the number of close neighbors each point has. The perplexity value has a complex effect on the resulting pictures. The original paper says, “The performance of SNE is fairly robust to changes in the perplexity, and typical values are between 5 and 50.” But the story is more nuanced than that. Getting the most from t-SNE may mean analyzing multiple plots with different perplexities.

In the Ramachandran plot, low dimensional projection of data along any one of the projections (*ϕ* or *ψ*), yields overlap of different conformations onto each other. We show this at the bottom and left of Fig. S2 (a) for projection along *ϕ* and *ψ*, respectively. PCA, the most common dimensional reduction method, fails to achieve clear separation and has a very low Silhouette score of 0.154 (see Fig. S2(b)). This is because PCA tries to linearly transform the data along an axis of maximal variation, which is the *ψ* axis in the Ramachandran plot and hence cannot capture the distinction between L-helix and other conformers. On the other hand, the t-SNE projections provide more faithful representations of the clusters. In Fig. S2 (c), we plot the t-SNE projections for a range of perplexity values. For a certain perplexity value (*Perp* =400), t-SNE clearly separates out the 4 sub-regions as in the original space with a much improved Silhouette score of 0.50. At low perplexities, t-SNE focuses on the local variations and tries to preserve the closest neighbors as much as possible in the original space. However, very low perplexity yields too many clusters with single or very few conformations per cluster, which nullifies the advantages of clustering in the first place. On the other hand, t-SNE essentially degrades to PCA at very high perplexities and leads to overcrowding as greater variations are tolerated at high perplexity scores. With perplexity as a tuneable parameter to balance the degree of local preservation on one hand and minimize the overcrowding on the other, t-SNE offers an exciting possibility to meaningfully cluster and visualize complex and heterogeneous high-dimensional IDPs datasets.

**FIG. S2:**
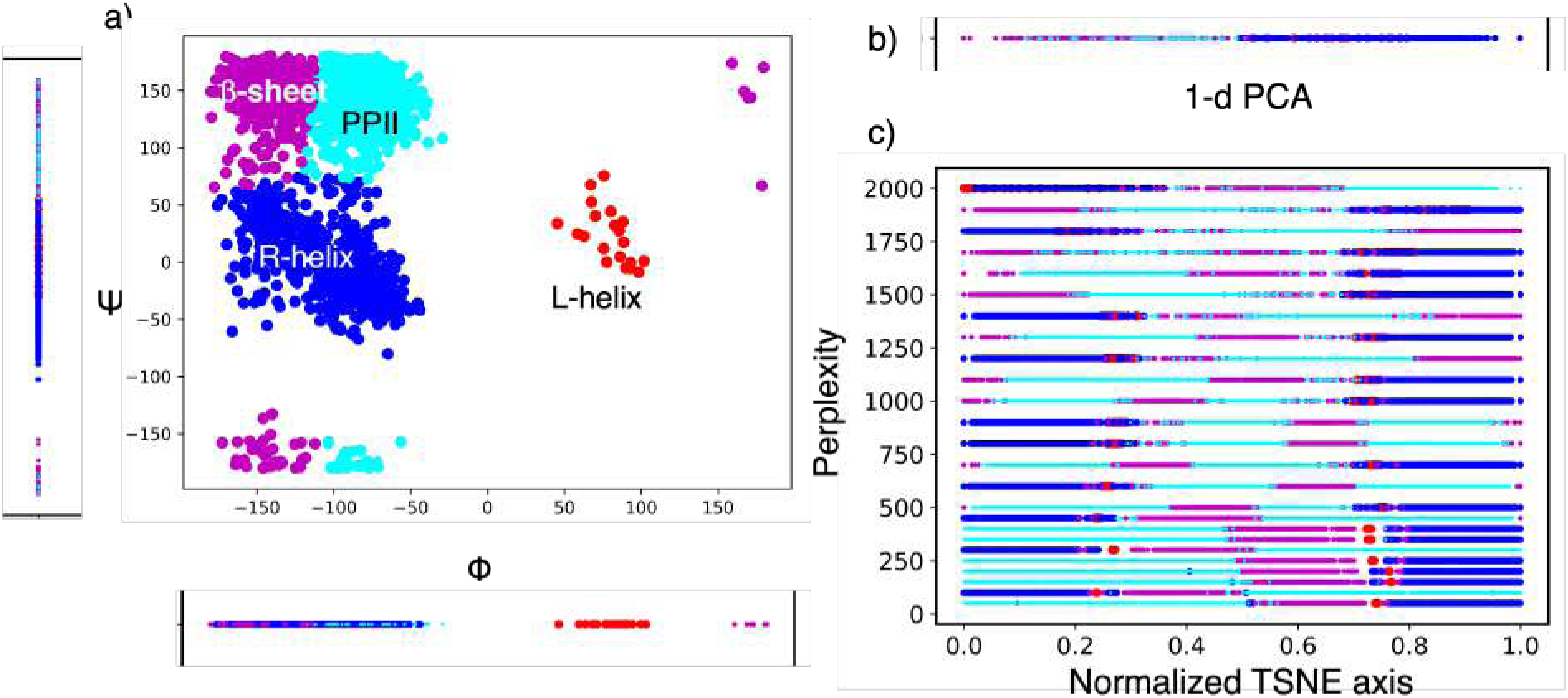
Dimensional reduction of Ramachandran map of alanine di-peptide: a) The 2D Ramachandran map color-coded by the subregions. Simple 1-dimensional projections along the X and Y axes are shown. Results of 1D transformation of RC-map using b) PCA and c) t-SNE with different perplexities.

### B Estimation of homogeneity within cluster

To quantify the homogeneity of conformations within a cluster, we first reorder the conformations based on the cluster indices and plot their pairwise similarity/distances. Fig. S3 and Fig. S4 show the results for the APO and the G5 bound A*β*42 ensemble, respectively. We also report the distance map before clustering for comparison. The conformational distance is measured based on the RMSD of inter-residue LJ energies. To further accentuate the homogeneity illustrations, we have also plotted the respective RMSD of Cartesian coordinates. Please note that for the clustering with t-SNE, only the RMSD of LJ energies was used. For the input maps (Fig S3a and S4a), we ordered the conformations sequentially in the X and Y axes. For the maps generated after clustering, the frames are sorted based on the cluster indices and placed from 0*^th^* cluster to N*^th^* cluster (S3-S4, b-d for both the upper and lower panel in Fig. S3 and Fig. S4, respectively). As indicated by the figures, the input distance maps of these ensembles show a certain level of conformational memory across the contiguous frames (the Red blocks/grids at the diagonal band) as the trajectories are generated from a history-dependent metadynamics approach. Nevertheless, the clustering obtained with sub-optimal parameters adversely affects even this intrinsically clustered data and several off-diagonal Red patches appear in the plot indicating either wrong groupings or broken clusters. On the other hand, with the optimal parameters, the algorithm yields better clustering. This can be seen clearly when we remove the input bias by shuffling the frames randomly and subjecting them to t-SNE. The resultant clustering on the shuffled data with optimal parameters indicates the proper grouping of conformations with diagonal Red blocks and no off-diagonal Red patches (Compare Figure S3 and Figure S5). Also the resultant cluster assignments with both unshuffled and shuffled data are consistent with a mutual information score of 0.96, indicating faithful clustering irrespective of the order of input, upon using optimal parameters.

**FIG. S3:**
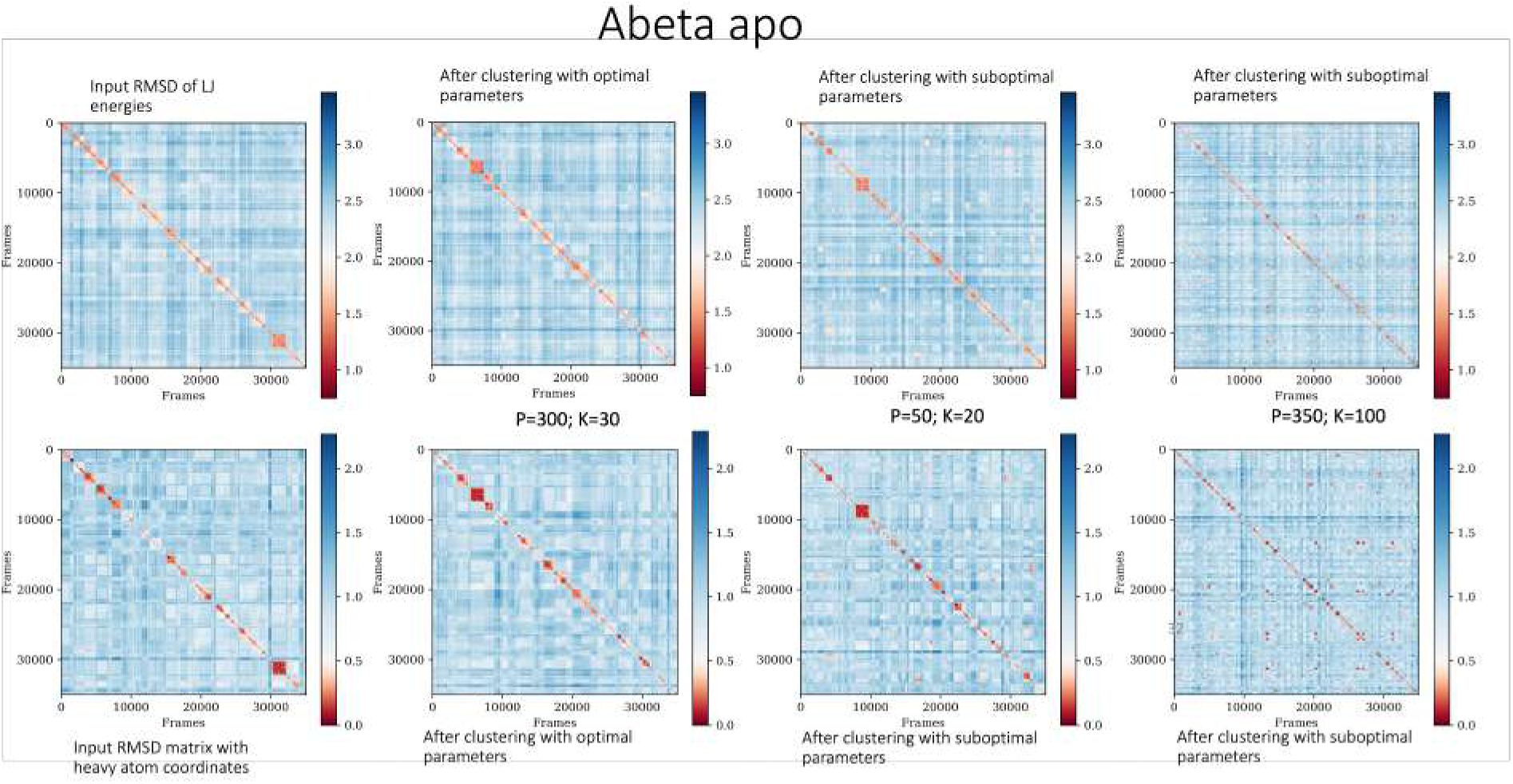
Pairwise RMSD map of apo A*β*42 conformations: Row-1 and Row-2 represents the RMSD map generated on the inter-residue LJ energies and on the heavy atom coordinates of conformations, respectively. The map generated on the raw trajectory (before clustering) is shown in a and e. The plots made after clustering is shown in b-d and f-h, where the frames are reordered according to the cluster indices (from the 0th cluster to the Nth cluster). The P and K values used for the clustering is indicated.

**FIG. S4:**
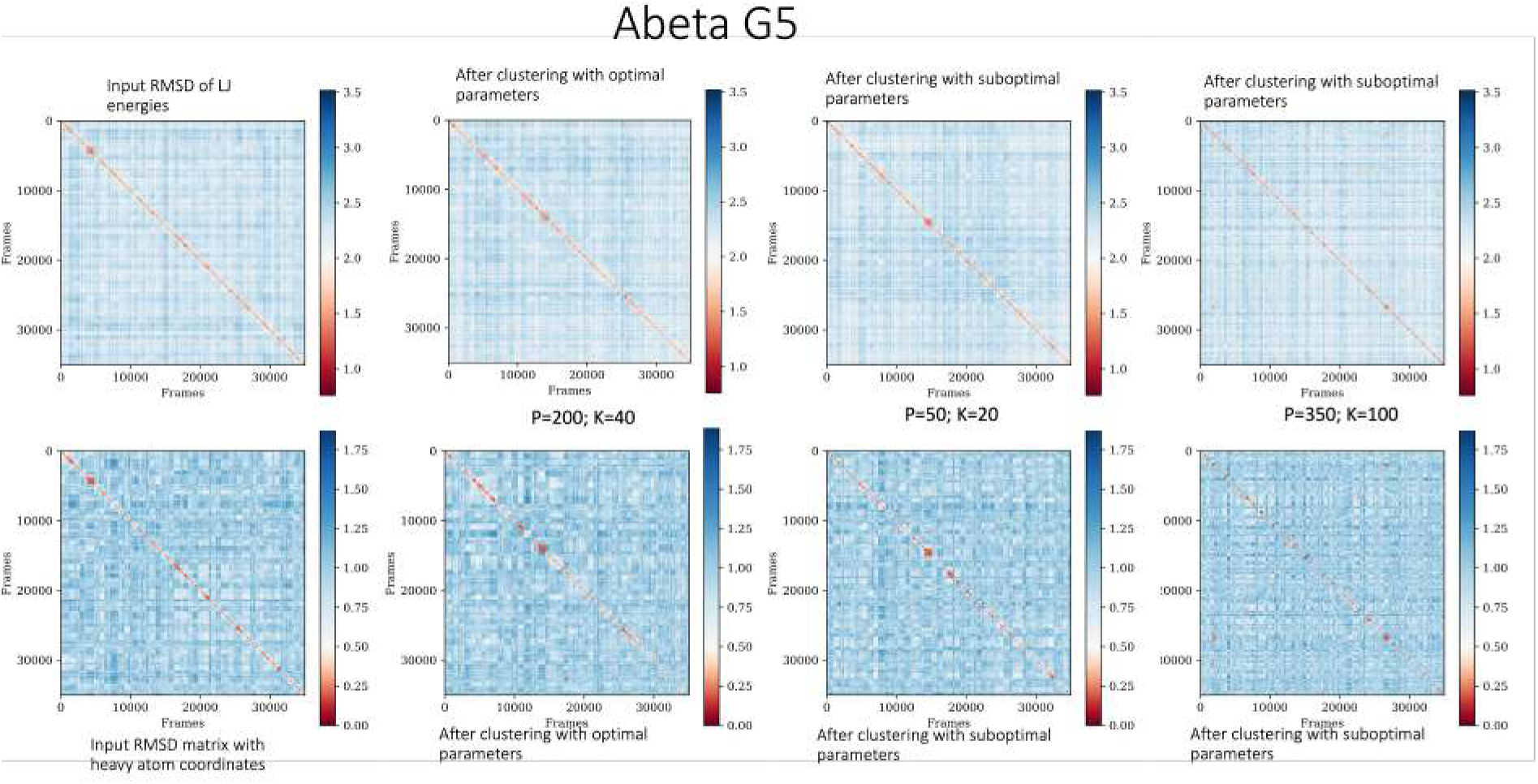
Pairwise RMSD map of G5-bound A*β*42 ensemble: Row-1 and Row-2 represents the RMSD map generated on the inter-residue LJ energies and on heavy atom coordinates of the conformations, respectively. The map generated on the raw trajectory (before clustering) is shown in a and e. The plots made after clustering is shown in b-d and f-h, where the frames are reordered according to the cluster indices (from the 0th cluster to the Nth cluster). The P and K values used for the clustering are indicated.

**FIG. S5:**
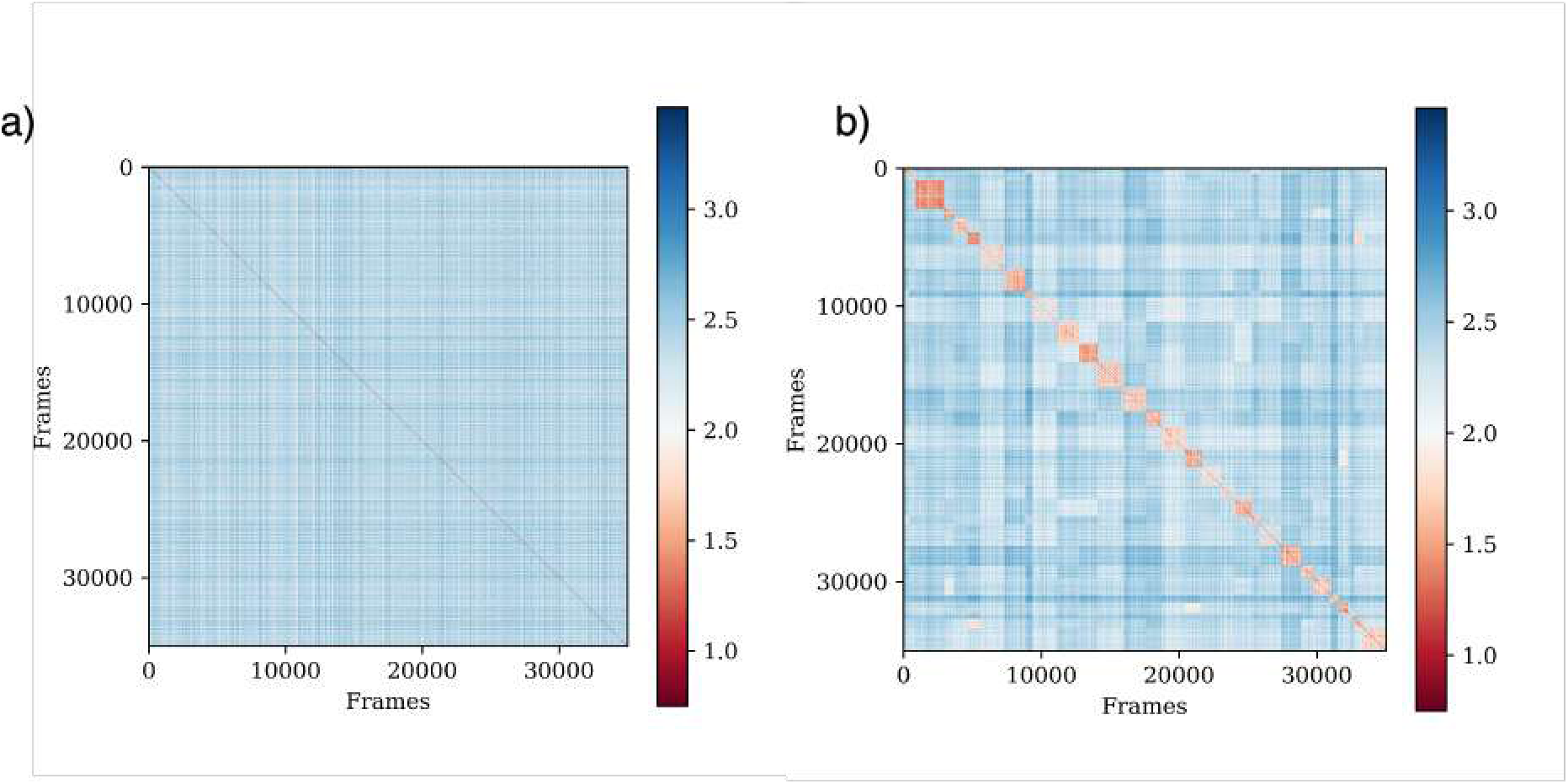
a) Pairwise RMSD map of apo A*β*42 conformations shuffled randomly. b) Pairwise RMSD map generated after clustering with optimal parameter set (P=300; K=30). The RMSD between conformations is calculated on the inter-residue LJ energies.

**FIG. S6:**
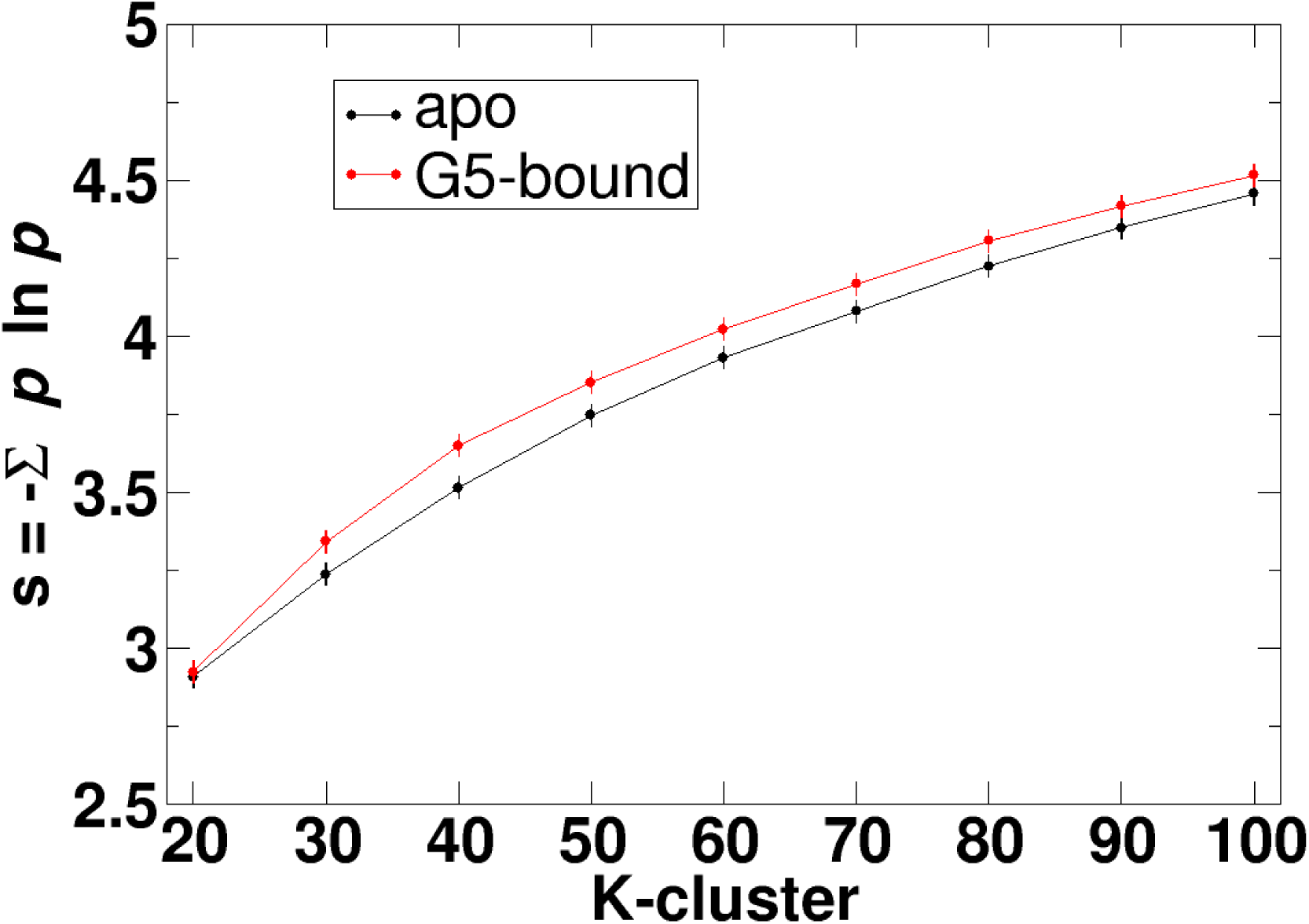
Estimation of Gibbs conformational entropy (*S* = −∑(*p_i_* ln *p_i_*)) in the apo and G5-bound A*beta*42 ensembles. *p_i_* is the fractional occupancy of each cluster, weighted by the metadynamics weights obtained from^4^.

**FIG. S7:**
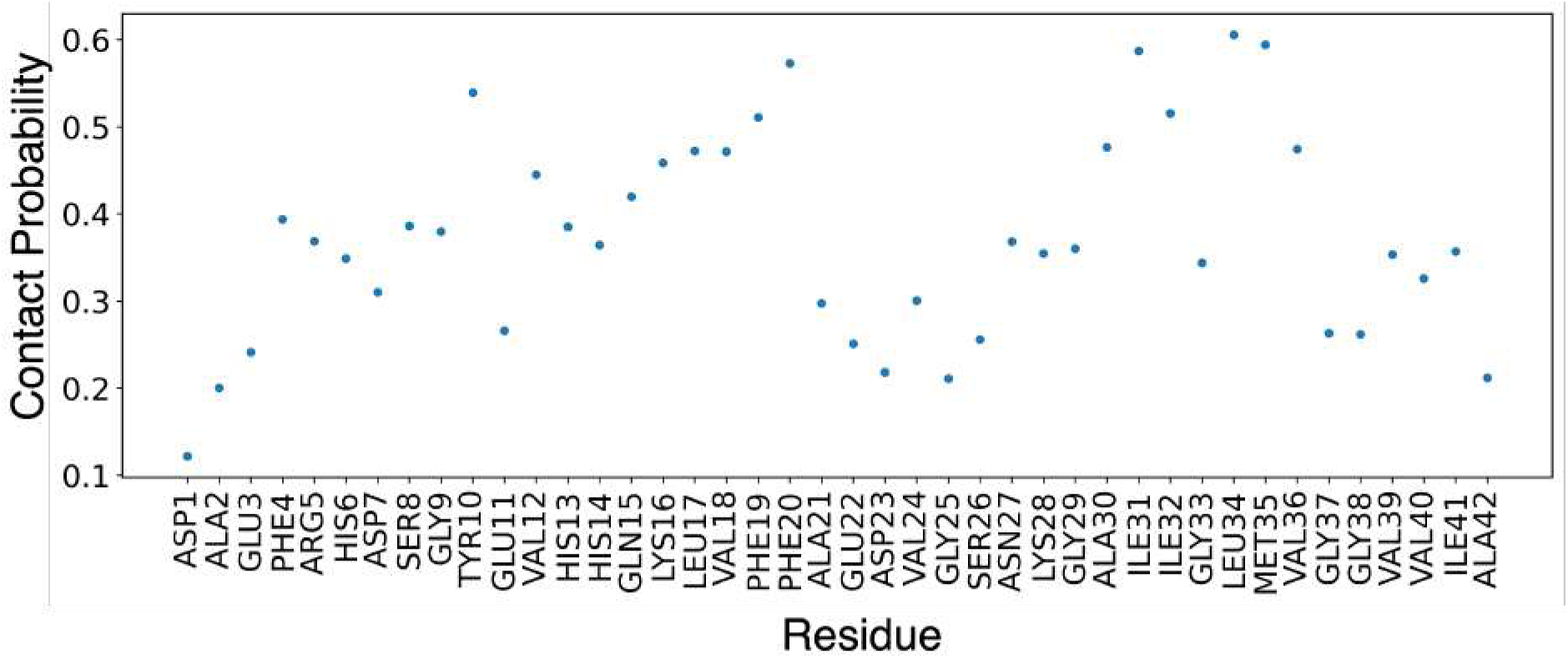
Residue-wise Propensity of contacts made by A*β*42 with G5 calculated from the total trajectory.

**FIG. S8:**
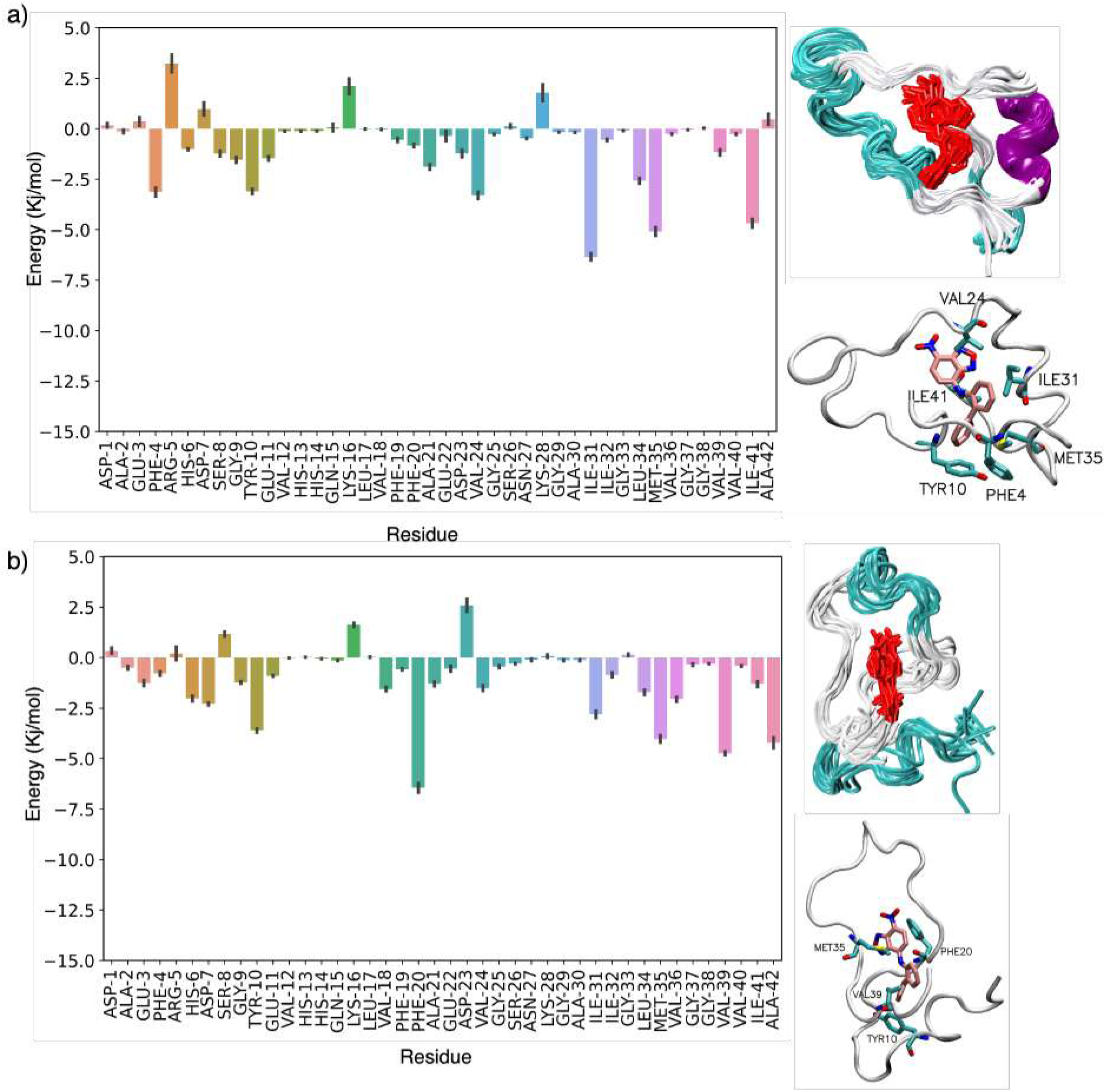
Residue-wise decomposition of total binding energy from the other two favorable bound geometries (cluster29 in (a) and cluster 30 in (b))

**FIG. S9:**
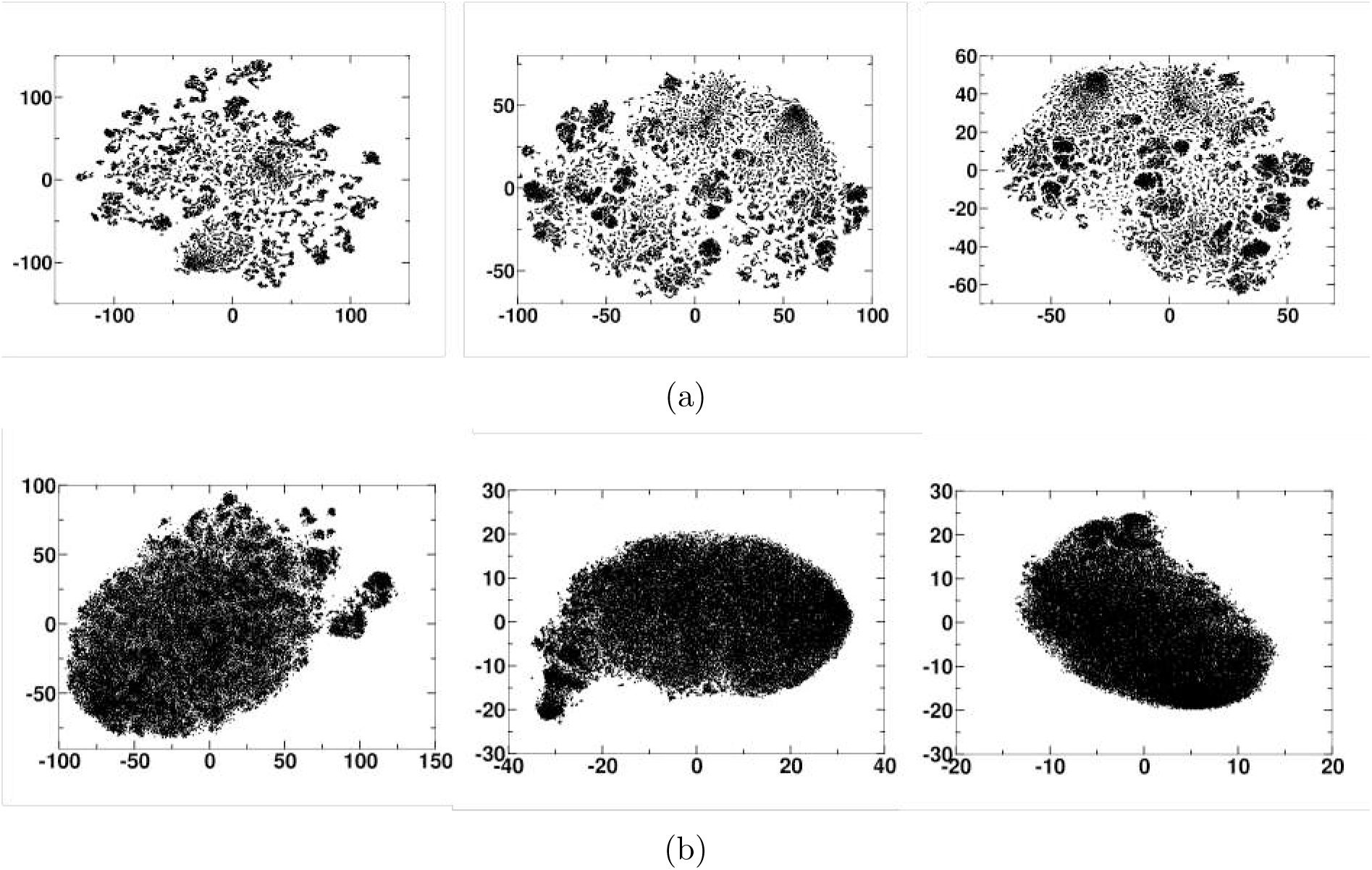
(a) Representative t-SNE maps generated with three different perplexity values (100, 1000 and 2000) for the full length apo *α*-synuclein ensemble (b) Representative t-SNE maps generated with three different perplexity values (100, 1000 and 2000) for the apo c-terminus *α*-synuclein ensemble

**FIG. S10:**
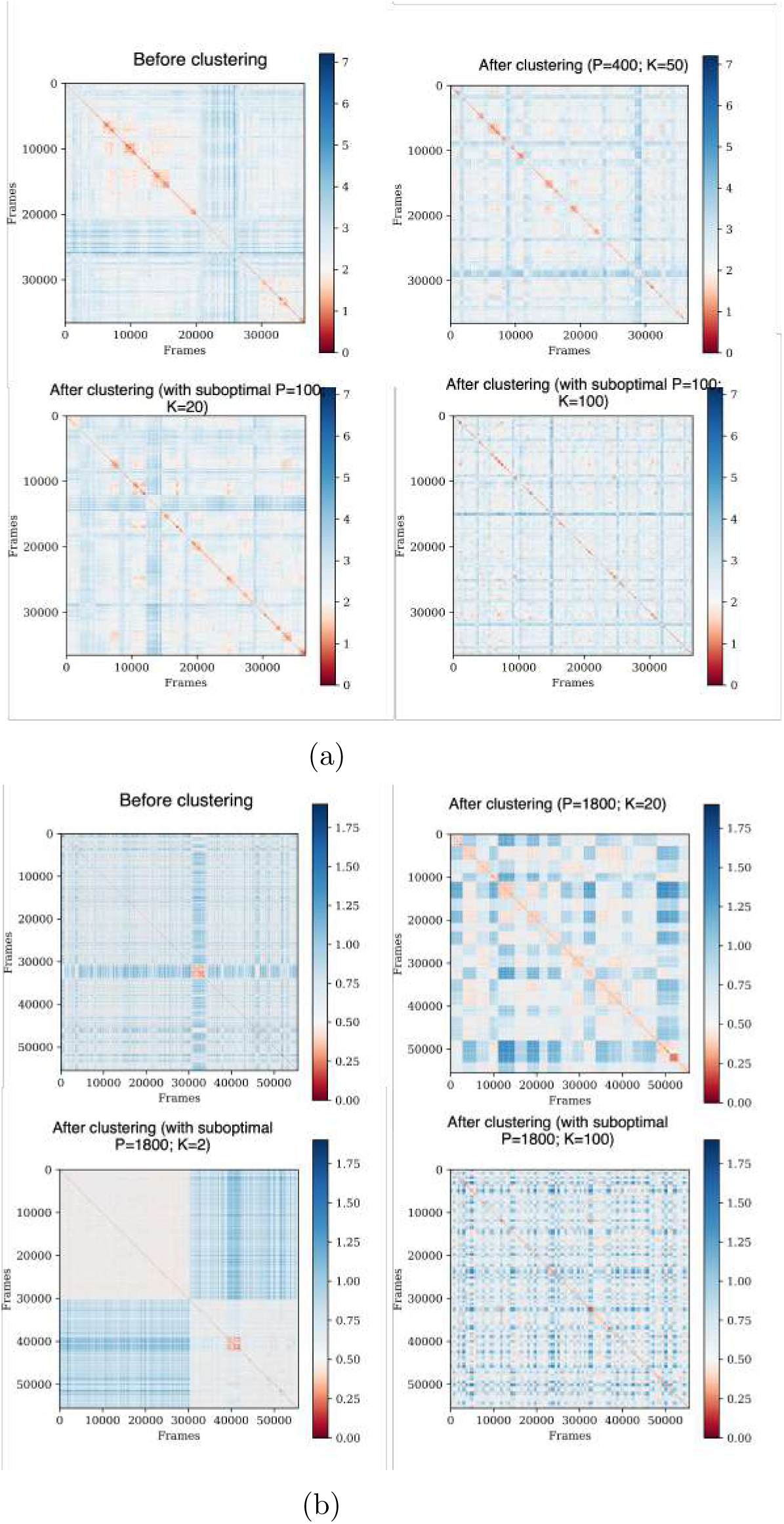
Pairwise RMSD map of (a) full-length apo and (b) c-terminus *α*S apo ensemble: The RMSD between each pair of conformation is measured based on the heavy atom coordinates. The map generated on the raw trajectory (before clustering) is shown in a. The plots made after clustering is shown in b-d, where the frames are reordered according to the cluster indices (from the 0th cluster to the Nth cluster). The P and K values used for the clustering are indicated.

**FIG. S11:**
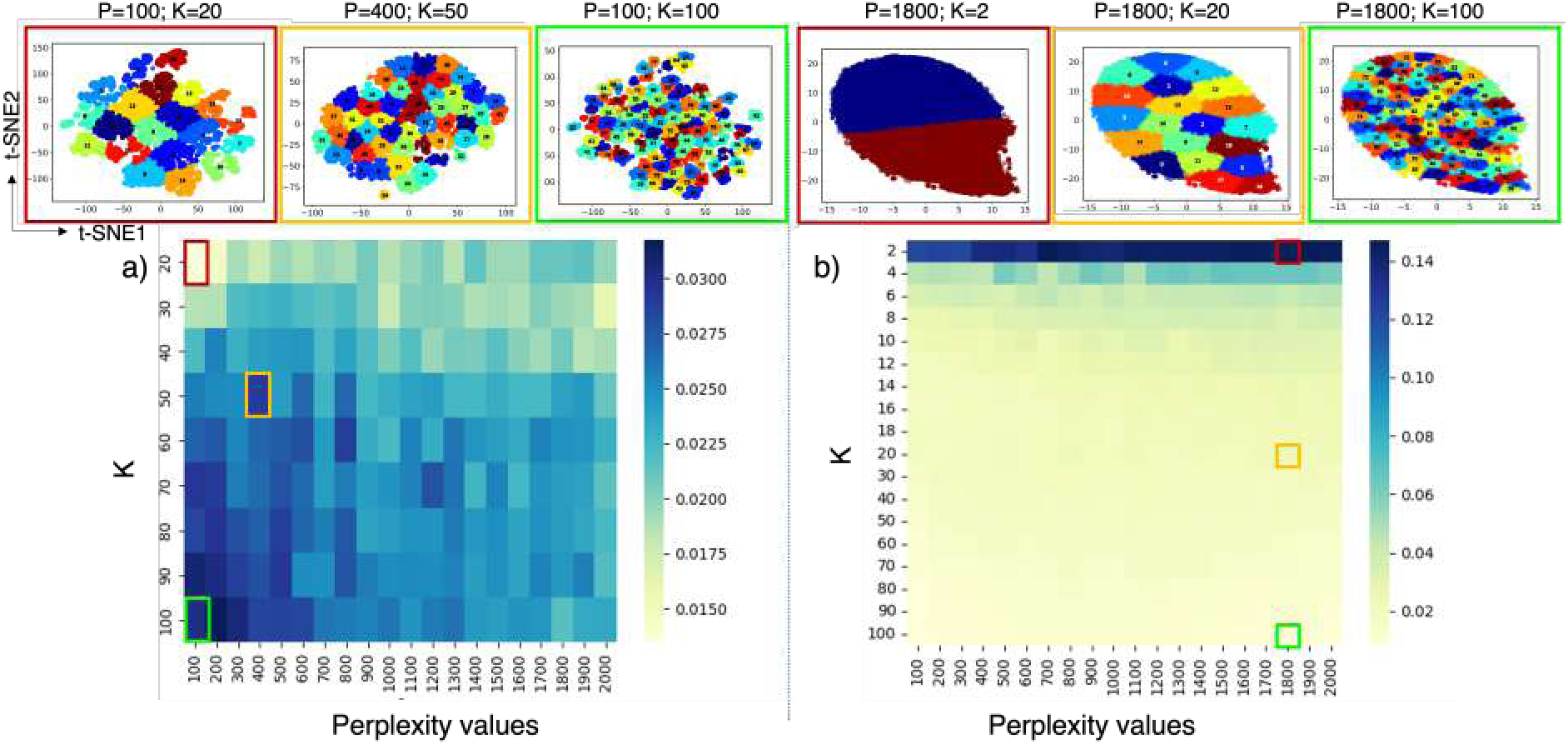
Hyperparameter optimization based on integrated Silhouette score for the (a) apo full-length and (b) apo c-terminal *α*-synuclein ensembles.

**FIG. S12:**
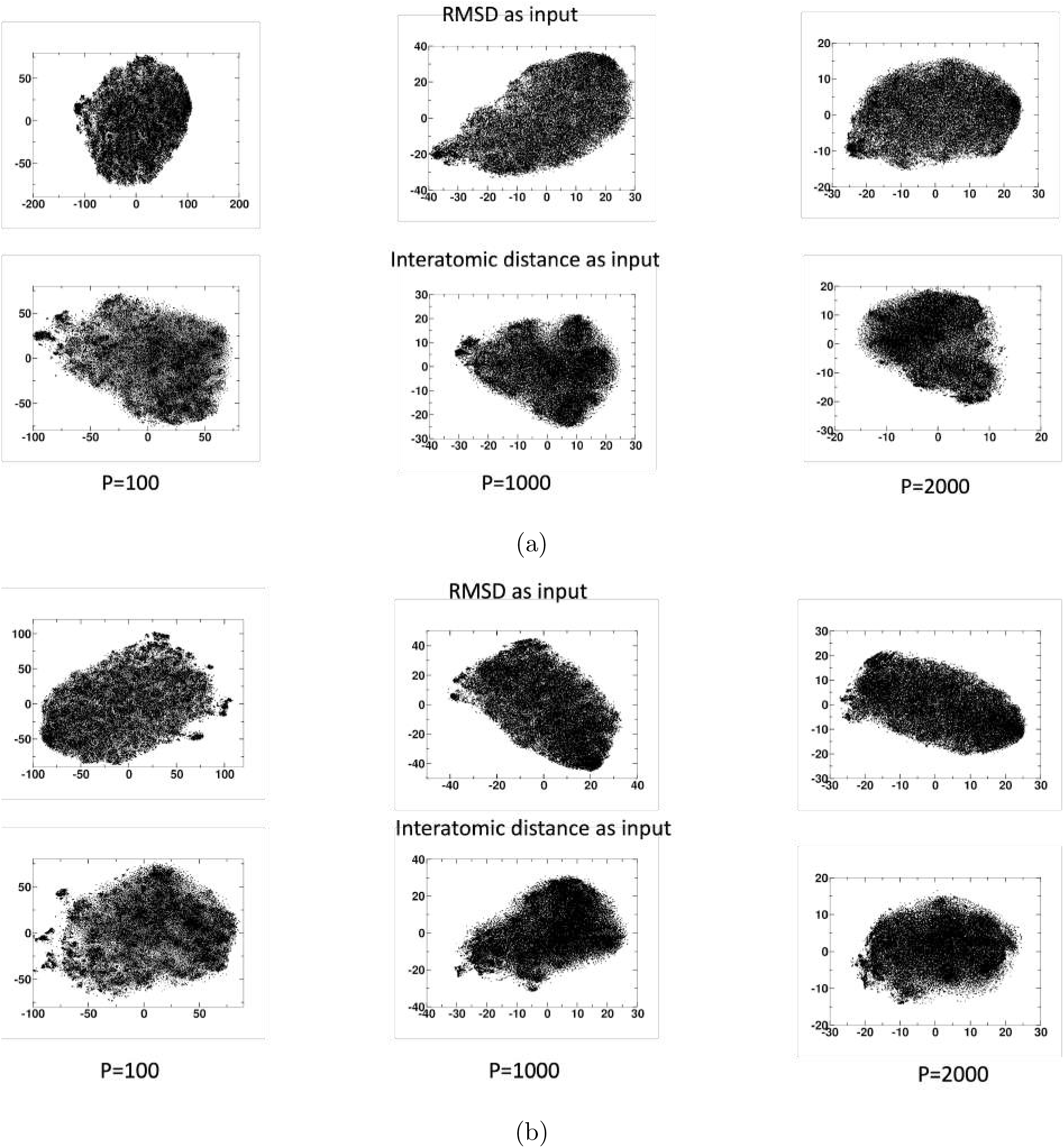
Representative t-SNE maps were generated with three different perplexity values (100, 1000, and 2000) for the (a) Fasudil-bound and (b) Lig47-bound c-terminus *α*-synuclein ensemble

**FIG. 13:**
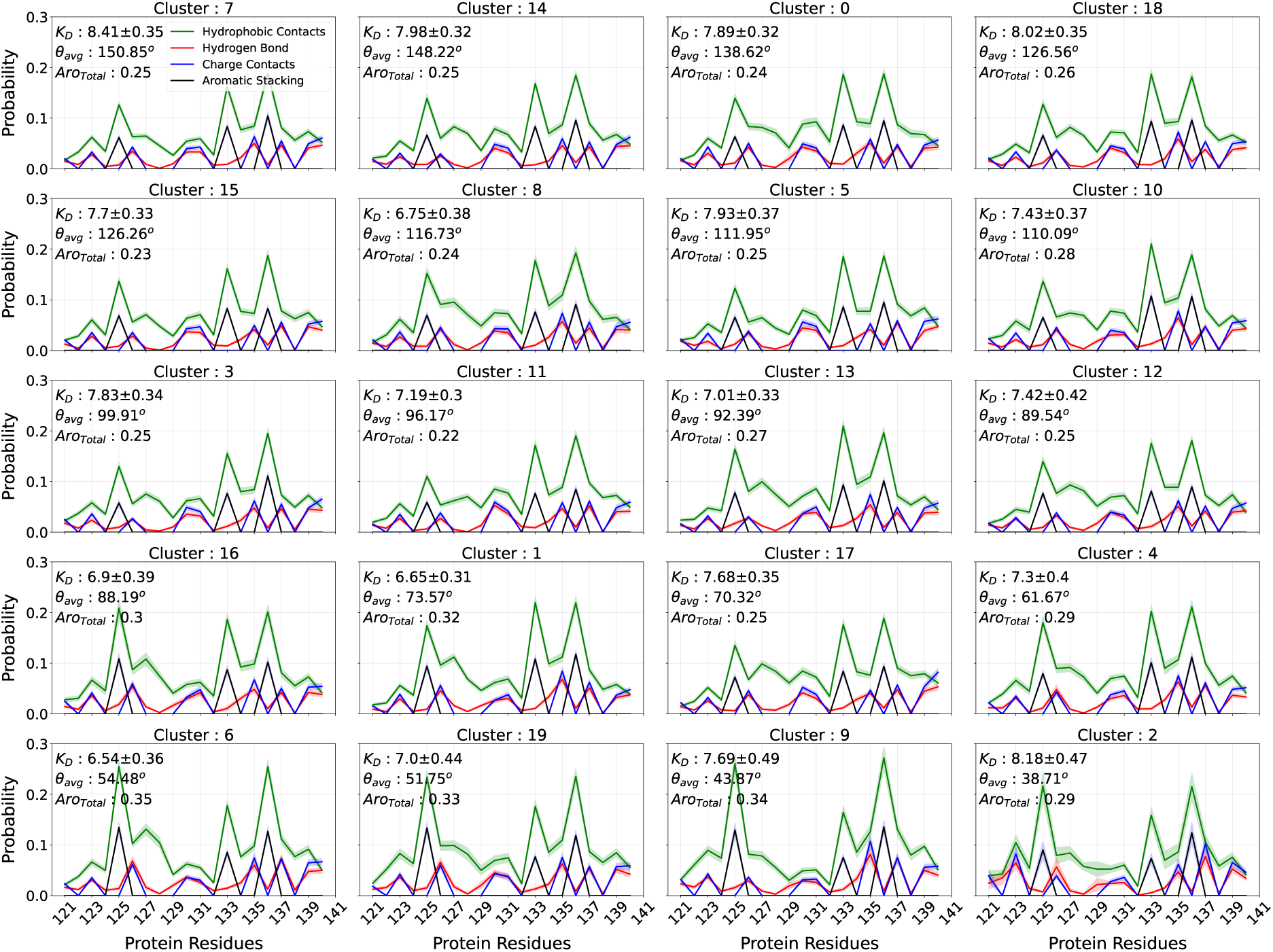
Fraction of specific inter-molecular interactions such as hydrophobic contact, aromatic stacking interaction, charge-charge contact and hydrogen bonding interaction between fasudil and each residue of *α*S C-term. The subplots are arranged based on descending order of average bend angle of clusters.

**FIG. 14:**
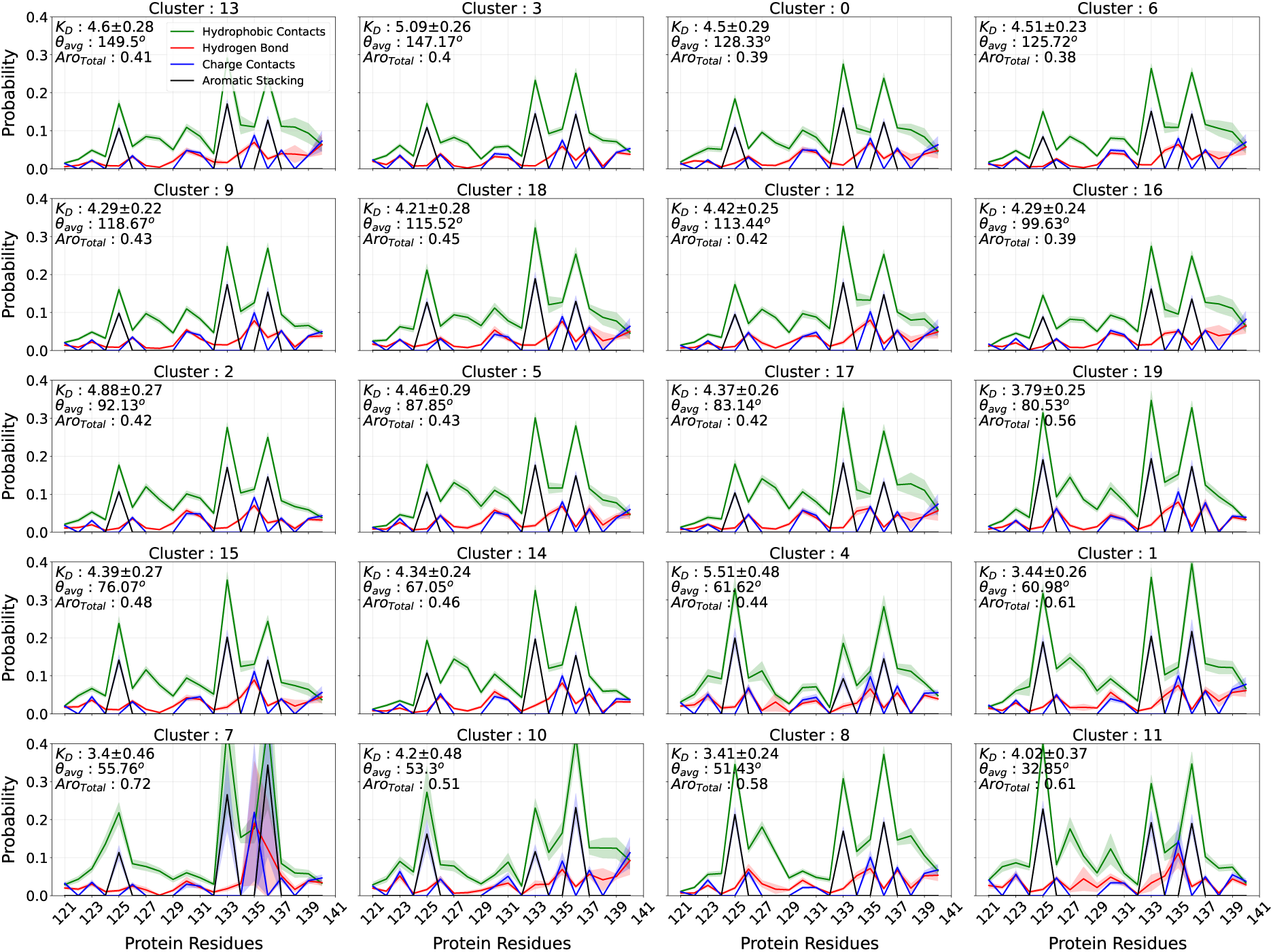
Fraction of specific inter-molecular interactions such as hydrophobic contact, aromatic stacking interaction, charge-charge contact and hydrogen bonding interaction between ligand-47 and each residue of αS C-term. The subplots are arranged based on descending order of average bend angle of clusters.

**FIG. 15:**
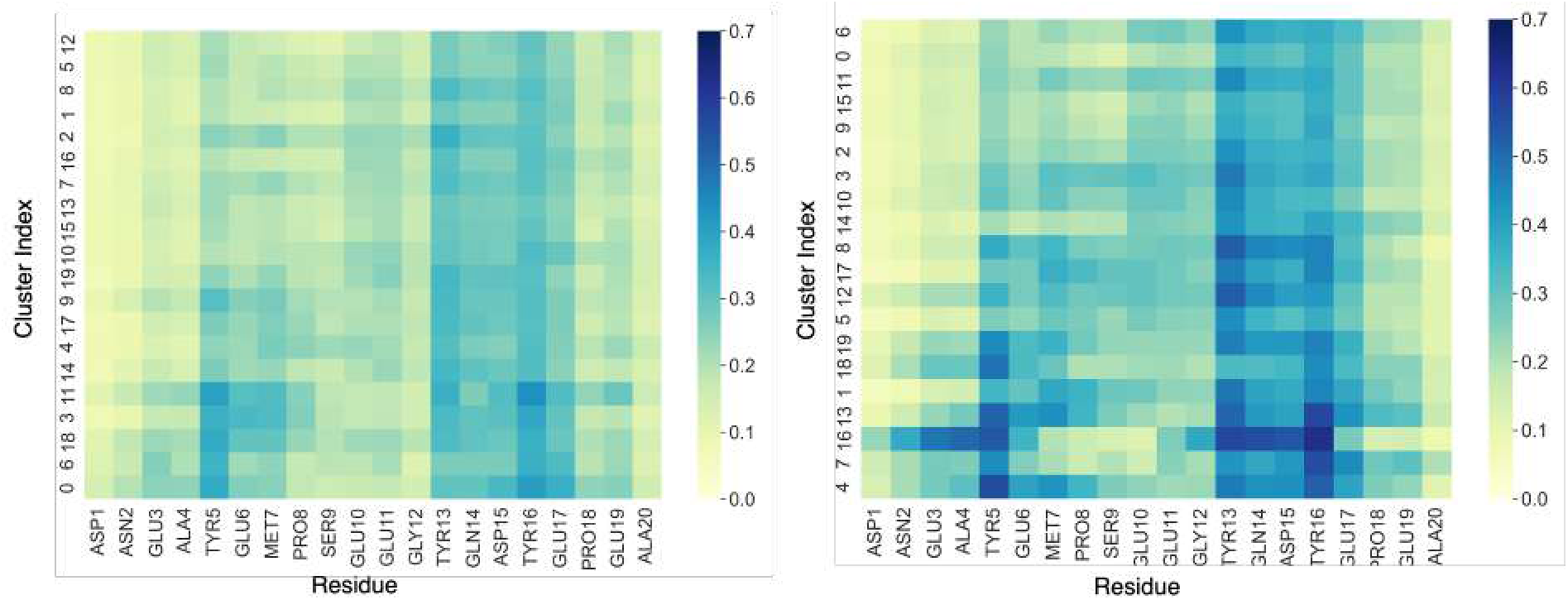
Per-residue inter molecular contact probabilities calculated after clustering the bound frames alone. The contacts between *α*S*_C−term_* and fasudil and *α*S*_C−term_* and ligand 47 observed in each cluster are shown in (a) and (b) respectively. The clusters are sorted in the decreasing order of bend angle. Note: The obtained clusters had similar bend profiles (data not shown).

**FIG. 16:**
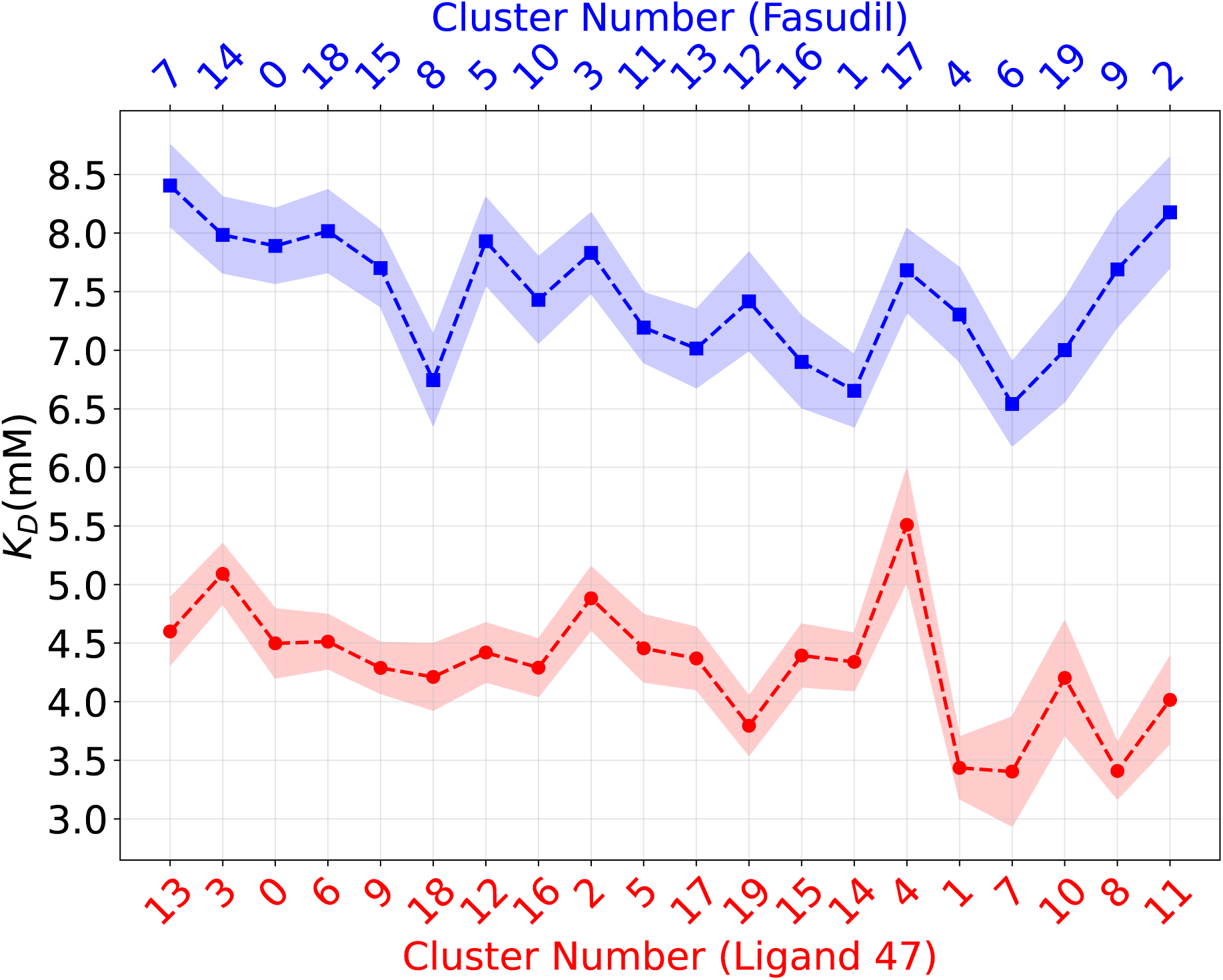
Estimated *KD* values of fasudil and ligand 47 across different clusters. The clusters are sorted in descending order of their average bend angle.

**FIG. 17:**
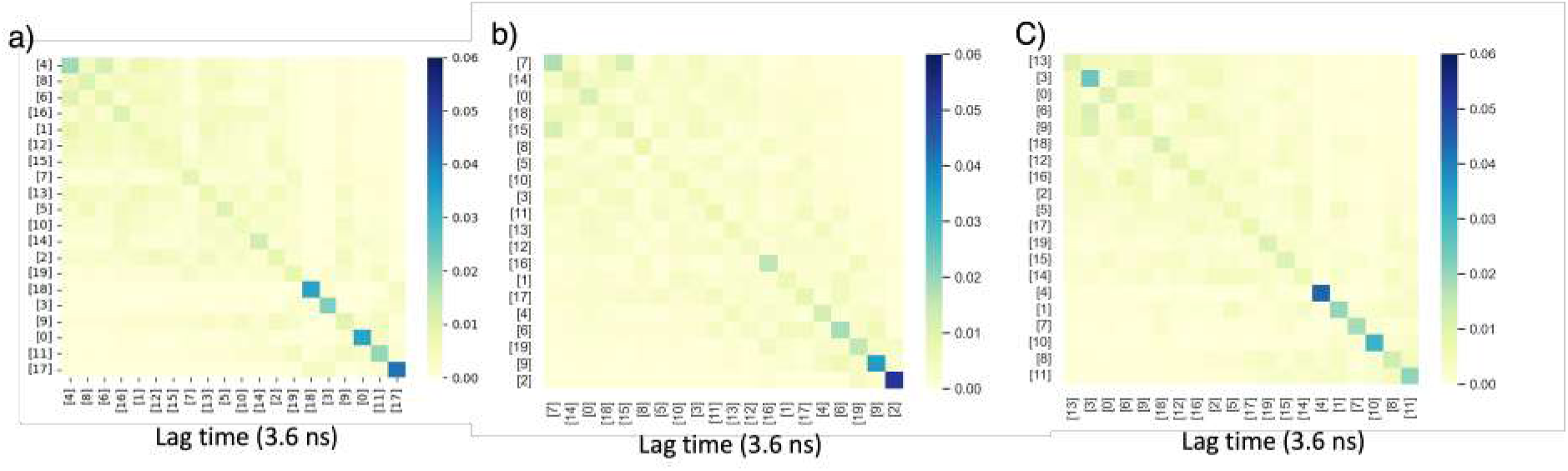
Cluster-wise transition probability of (a) apo, (b) fasudil bound and (c) ligand 47 bound *α*-synuclein C-terminal peptide conformations at a lag time of 3.6 ns.

**FIG. 18:**
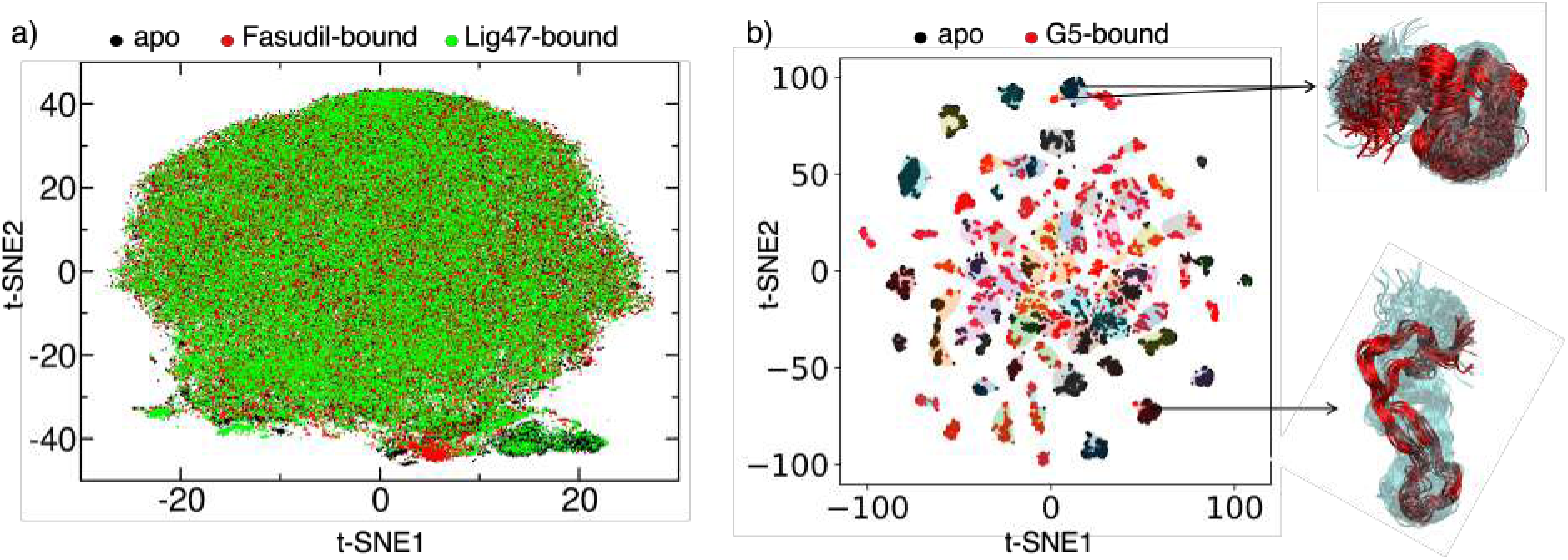
a) t-SNE map of collated ensembles of *α*-synuclein C-terminal peptide simulated in the presence and absence of Fasudil or ligand-47. The reduced coordinates of apo, Fasudil-bound, and Ligand-47 bound data are color coded in Black, Red, and Green respectively. b) t-SNE map of collated ensembles of A*β*42 protein simulated in the presence (red dots) and absence of G5 (black dots). The map shows mostly non-overlapping clusters between the two ensembles except for a very few clusters that consist of conformations from both apo and bound ensembles. These similar and overlapped conformations from the two ensembles are rendered and superposed in the inset with cyan indicating apo and red indicating G5-bound A*β*42 ensembles. We also mapped the conformational clusters that showed the highest binding affinity with G5 molecules (cluster numbers 14, 29, and 30) as predicted from Fig. 3 in the main text and indicate them on the collated map with blue arrows. It is evident that these high-affinity conformations from the G5-bound ensemble do not have closely related conformers in the APO state.

**TABLE S1:** Mean Silhouette score for the apo Aβ42 ensemble calculated based on the distances in the low S_ld_ and high dimensional space S_hd_. S_hd_ values are given in the bracket

|  | 20 | 30 | 40 | 50 | 60 | 70 | 80 | 90 | 100 |
| --- | --- | --- | --- | --- | --- | --- | --- | --- | --- |
| 50 | 0.424<br>(0.087) | 0.498<br>(0.148) | 0.546<br>(0.171) | 0.576<br>(0.157) | 0.583<br>(0.139) | 0.598<br>(0.13) | 0.593<br>(0.119) | 0.589<br>(0.109) | 0.583<br>(0.108) |
| 100 | 0.505<br>(0.141) | 0.586<br>(0.177) | 0.64<br>(0.189) | 0.642<br>(0.163) | 0.633<br>(0.139) | 0.611<br>(0.125) | 0.598<br>(0.116) | 0.584<br>(0.11) | 0.582<br>(0.107) |
| 150 | 0.543<br>(0.15) | 0.626<br>(0.211) | 0.686<br>(0.194) | 0.658<br>(0.166) | 0.639<br>(0.141) | 0.625<br>(0.13) | 0.598<br>(0.119) | 0.581<br>(0.112) | 0.574<br>(0.109) |
| 200 | 0.579<br>(0.166) | 0.666<br>(0.197) | 0.711<br>(0.195) | 0.665<br>(0.164) | 0.62<br>(0.133) | 0.616<br>(0.13) | 0.599<br>(0.124) | 0.581<br>(0.116) | 0.558<br>(0.107) |
| 250 | 0.617<br>(0.174) | 0.689<br>(0.212) | 0.706<br>(0.193) | 0.658<br>(0.161) | 0.653<br>(0.151) | 0.605<br>(0.129) | 0.588<br>(0.121) | 0.572<br>(0.116) | 0.553<br>(0.108) |
| 300 | 0.638<br>(0.178) | 0.709<br>(0.211) | 0.712<br>(0.193) | 0.66<br>(0.164) | 0.617<br>(0.144) | 0.594<br>(0.132) | 0.569<br>(0.117) | 0.57<br>(0.116) | 0.545<br>(0.107) |
| 350 | 0.662<br>(0.188) | 0.718<br>(0.205) | 0.699<br>(0.192) | 0.637<br>(0.157) | 0.616<br>(0.141) | 0.593<br>(0.133) | 0.57<br>(0.119) | 0.563<br>(0.114) | 0.543<br>(0.105) |

**TABLE S2:** Mean Silhouette score for the G5-bound A*β*42 ensemble calculated based on the distances in the low S*_ld_* and high dimensional space S*_hd_*. S*_hd_* values are given in the bracket

|  | 20 | 30 | 40 | 50 | 60 | 70 | 80 | 90 | 100 |
| --- | --- | --- | --- | --- | --- | --- | --- | --- | --- |
| 50 | 0.418<br>(0.057) | 0.464<br>(0.097) | 0.513<br>(0.131) | 0.543<br>(0.132) | 0.554<br>(0.13) | 0.577<br>(0.127) | 0.578<br>(0.127) | 0.567<br>(0.116) | 0.584<br>(0.123) |
| 100 | 0.473<br>(0.072) | 0.555<br>(0.118) | 0.612<br>(0.149) | 0.63<br>(0.145) | 0.616<br>(0.141) | 0.618<br>(0.139) | 0.581<br>(0.122) | 0.57<br>(0.121) | 0.556<br>(0.119) |
| 150 | 0.487<br>(0.082) | 0.587<br>(0.126) | 0.652<br>(0.148) | 0.645<br>(0.145) | 0.627<br>(0.14) | 0.604<br>(0.14) | 0.577<br>(0.128) | 0.553<br>(0.124) | 0.554<br>(0.128) |
| 200 | 0.495<br>(0.081) | 0.605<br>(0.129) | 0.667<br>(0.149) | 0.66<br>(0.149) | 0.635<br>(0.146) | 0.598<br>(0.137) | 0.573<br>(0.137) | 0.518<br>(0.12) | 0.509<br>(0.121) |
| 250 | 0.508<br>(0.089) | 0.615<br>(0.13) | 0.662<br>(0.149) | 0.643<br>(0.148) | 0.61<br>(0.146) | 0.604<br>(0.144) | 0.545<br>(0.128) | 0.516<br>(0.125) | 0.491<br>(0.121) |
| 300 | 0.517<br>(0.094) | 0.596<br>(0.126) | 0.649<br>(0.151) | 0.63<br>(0.151) | 0.604<br>(0.147) | 0.531<br>(0.131) | 0.527<br>(0.127) | 0.51<br>(0.127) | 0.474<br>(0.119) |
| 350 | 0.509<br>(0.093) | 0.591<br>(0.129) | 0.633<br>(0.146) | 0.622<br>(0.15) | 0.592<br>(0.151) | 0.522<br>(0.124) | 0.492<br>(0.125) | 0.489<br>(0.124) | 0.48<br>(0.12) |

**TABLE S3:** Consistent local clustering in the apo Aβ42 trajectory upon different runs with different random initializations. We have noted that there is neither change in the choice of optimal perplexity and K values, nor the clustering pattern (measured by normalized mutual information score) upon rerunning.

| S.No | $S_{ld}$ | $S_{hd}$ | Normalized mutual information |
| --- | --- | --- | --- |
| 1 | 0.7086 | 0.2105 | 0.9755 |
| 2 | 0.7158 | 0.2102 |  |

